# The lysine side chain’s swing motion in AapA1 toxin promotes water transport through transmembrane pores

**DOI:** 10.1101/2025.03.24.644837

**Authors:** Zhihong Shi, Lei Liu, Mingqiong Tong, Yaqing Fang, Dongying Yang, Liling Zhao, Zanxia Cao

**Affiliations:** Dezhou University

**Keywords:** toxin-antitoxin system, AapA1-membrane, multimeric toxin protein structural characteristics, pore formation, molecular dynamics simulation

## Abstract

The rising antibiotic resistance of *Helicobacter pylori* and the emergence of multi-drug-resistant strains pose significant challenges to its eradication. Understanding the molecular mechanisms driving this resistance is critical. This study investigates how AapA1, a toxin protein, may inhibit *H. pylori* growth and cause morphological changes by forming transmembrane pores in its inner membrane. Using experimental data and molecular dynamics simulations, we analyzed the pore structure and its formation mechanism. Results show that AapA1 forms pores on liposomes and exists stably in the membrane as α-helices through different multimers, creating hydrophilic pores of varying sizes. Dynamic oscillation of K16/23 side chains in the membrane affects water transport through these pores, and restricting their movement significantly reduces transport. AapA1 monomers adopt three distinct conformations, with the K16/23 residues showing an up-down swing motion, potentially facilitating accelerated transport of substances. This study provides a new perspective on the dynamic structure–function relationship of AapA1-membrane interaction and highlights the role of positively charged residues in AapA1. Understanding the multimeric AapA1-membrane structure and the mechanism underlying AapA1-induced transmembrane pores is particularly crucial to develop new drugs and treatment options to reduce drug resistance emergence and combat any new emergence development.

## Introduction

*Helicobacter pylori*, a common human gastrointestinal pathogen, presents substantial challenges in its effective elimination from the host through the innate immune system ^1, 2^. Following external antibiotic threat perception, *H. pylori* cells undergo morphological changes and enter a dormant state, leading to increased antibiotic resistance and further complicating effective eradication measures ^3, 4^. However, the mechanisms through which *H. pylori* cells develop antibiotic resistance and show immune evasion remain poorly understood.

In *H. pylori*, the expression of the toxin protein AapA1 belonging to the type I toxin-antitoxin (TA) system ^5^ *aapA1*/IsoA1 ^6, 7^ induces the rapid morphological transformation of the cells from spiral shape to coccoid shape, resulting in growth inhibition and affecting the infectivity of *H. pylori* ^8^. In the dormant state, *H. pylori* cells exhibit a certain degree of clinical resistance to the combination of conventional therapy and vitamin D3 treatment ^9^. The growth inhibition of *H. pylori* is related to the structural function of AapA1 on the cell membrane. The findings of plasmon-waveguide resonance (PWR) and nuclear magnetic resonance (NMR) suggest that AapA1 targets the inner membrane of bacterial cells and inserts vertically into the membrane with a stable transmembrane α-helix conformation (PDB Id: 6GIG) ^10^. The growth inhibition of *H. pylori* and its morphological changes may be related to the oligomerized AapA1-induced formation of transmembrane pores with a refined architecture in the membrane.

Transmembrane pore formation induced by proteins/peptides is a common phenomenon in biological systems ^11, 12^. Transmembrane pores provide direct pathways for the entry and exit of substances in cells and can also cause cell death. These pores play a critical role in various biological functions vital for the maintenance of complex organisms; moreover, disruptions in these processes can lead to several diseases. A wide variety of proteins can form pores and perform different functions in the cell membrane. For example, gasdermins are used to induce pyroptosis; mixed lineage kinase domain-like proteins are the essential elements of necroptosis; pore-forming toxins ^13–15^ are one of the most potent virulence factors secreted by bacteria during infection, and they disrupt the cell membrane by forming stable pores; and aquaporins are pore-forming proteins that facilitate water diffusion through the cell membrane ^16–18^. These proteins from different organisms share a common mechanism of action, wherein the protein transmembrane region inserts into the membrane lipid bilayer to form water-filled pores; however, the pores formed by these proteins differ in structure, properties, and size. Over the past several years, some studies have investigated the structure, properties, and size of the pores at the atomic level ^19–23^. Interestingly, pore-forming motifs of these proteins share very low sequence similarity with each other; consequently, the structure of other pore-forming proteins cannot be used as a guide for determining AapA1 structure ^11, 14^. Hence, it is critical to study the structural characteristics and pore-forming mechanisms of AapA1 in the membrane.

Enhanced comprehension of AapA1-membrane conformation dynamics and the mechanisms underlying pore formation on the cell membrane could provide a basis to better understand bacterial drug resistance mechanisms and develop new drug targets. Various techniques, including fluorescence spectroscopy, fluorescence dequenching, circular dichroism, western blotting assay, zeta potential determination, solid-state NMR, PWR, cryo-electron microscopy, and molecular dynamics (MD) simulations, have been used for studying interactions between bacterial type I toxin proteins and the cell membrane ^5, 10, 24–27^. For instance, a series of experiments and MD simulations conducted in the past decade revealed that TisB is a type I toxin protein inserted vertically into the membrane as tetrameric bundles that decrease proton motive force and ATP levels ^25, 28–32^. AapA1 targets the inner cell membrane, and its monomers exist as transmembrane helices ^8, 10, 33^; however, the multimeric AapA1-membrane structure and the mechanism of action of multimeric AapA1 on the membrane remain unknown. AapA1 may adopt different multimeric structural states in the membrane, which implies that AapA1 may potentially form diverse peptide-membrane structures that can affect the pore-forming function. However, by utilizing the existing experimental methods, it is difficult to visualize the multimer structure and dynamics of AapA1 on the cell membrane.

Understanding the dynamic structural state of AapA1 in the bilayer membrane is crucial to elucidate its functions. MD simulation in conjunction with an enhanced sampling method is an effective tool to reveal protein-membrane interactions at the atomic level and strengthen experimental results ^16, 17, 20, 22, 34–38^. These simulations typically use techniques such as umbrella sampling, replica-exchange umbrella sampling, or various metadynamics methods to determine the free energy surface of the protein/peptide-cell membrane system.

Based on the aforementioned considerations, the present study aimed to investigate the critical forms of AapA1 by using MD simulations for multimeric AapA1-containing membrane systems in conjunction with bacterial cell membrane experiments involving liposomes. We found that AapA1, ranging from the tetrameric to nonameric form, can stably exist in the membrane and produce hydrophilic pores. The K16/K23 side chains of AapA1 multimer oscillate within the cell membrane to increase cell membrane perturbation and improve water permeability. The residues K16 and K23 of AapA1 may serve as a novel therapeutic target for the clinical management of gastric *H. pylori* by inhibiting the mechanisms involved in the transition to a drug-resistant bacterial phenotype.

## Materials and Methods

### 1. Experimental methods

#### Compliance with Ethics Requirements

This study was conducted in accordance with the Declaration of Helsinki and approved by the Ethics Committee of Dezhou University (approval number: DWLL202403). Informed consent was obtained from all participants, and their confidentiality was maintained throughout the research. Participants were informed of their right to withdraw at any time without consequences. All procedures were designed to minimize risks and ensure participant well-being.

#### Peptide synthesis

The 30-amino-acid peptides (AapA1: NH2-MATKHGKNSWKTLYLKISFLGCKVVVLLKR-COOH, AapA1-16/23: NH2-MATKHGKNSWKTLYL**A**ISFLGC**A**VVVLLKR-COOH) were synthesized by standard solid-phase synthetic peptide chemistry and purified by high-performance liquid chromatography (Xian Peptide Biological Technology Co., Ltd.). **See supporting information for MS analysis report**. Because AapA1 and AapA1-16/23 are insoluble in water, they were dissolved in dimethyl sulfoxide (DMSO) and diluted with PBS to achieve the final concentration.

#### Hemolytic activity

Hemolytic activity was determined as described previously ^39^. AapA1 peptides at different concentrations were mixed with human erythrocytes in equal volumes and incubated for 30 min at 37 °C; saline and 0.5% Triton X-100 were used as negative and positive controls, respectively. After centrifugation at 2000 rpm for 5 min, the absorption was measured at 540 nm by using an Infinite 200 PRO microplate reader (Tecan, Männedorf, Switzerland). Hemolytic activity was calculated as follows: (experimental group – negative control) / (positive control – negative control) × 100%

#### Liposome preparation

Liposomes were prepared as described previously with minor modifications ^40^. Palmitoyloleoyl phosphatidylethanolamine (POPE) and palmitoyloleoyl phosphatidylglycerol (POPG) were purchased from Avanti Polar Lipids, Inc. The liposomes were prepared with the following composition ratio: 3:1 mole percent POPE: POPG. Briefly, the lipids in stock solutions prepared in CHCl_3_ were mixed at the desired molar ratio. The solvent was removed in a rotary evaporator. The lipid mixtures were then dried onto a film. Lipid films were then hydrated with 50 mM MES (pH 7.5) for 2 h at 37 °C. To prepare fluorescent-labeled liposomes, 50 mM MES solution (pH 7.5) containing 100 mM calcein was used as the hydration buffer. Liposomes were then generated by 10 freeze-thaw cycles in liquid nitrogen and 50 °C water bath. Large unilamellar vesicles (LUVs) were formed by extrusion of the generated liposomes through polycarbonate filters with 100 nm pore size (Avanti Polar Lipids, Inc.). Free lipids and fluorescent dyes were removed by centrifugation using a 100-kDa ultrafiltration tube. To prepare vitamin D3-enriched liposomes, 50 mM MES solution (pH 7.5) containing 10 μM vitamin D3 was used as the resuspension buffer. The prepared liposomes containing fluorescent lipids were stored at 4 °C in the dark and used within 2 days.

#### Nanoparticle tracking analysis (NTA)

The enriched liposomes were diluted in PBS and analyzed for particle size and concentration by using a nanoparticle tracking analyzer (ZetaView PMX 110; Particle Metrix, Meerbusch, Germany), and the data were analyzed with ZetaView 8.04.02 SP2 software.

#### Transmission electron microscopy (TEM)

The experiments were based on a previous report ^41^. Enriched liposomes were directly attached to Ni grids for 10 min before experimental treatment. The liposomes were stained with 2% uranyl acetate for 3 min before observation by using a transmission electron microscope (JEM-1400 plus) at 100 kV.

#### Calcein release assay

Calcein-loaded liposomes were mixed with peptides at different concentrations and incubated for 10 min at 37 °C; 0.5% DMSO and 0.5% Triton X-100 were used as negative and positive controls, respectively. Calcein fluorescence signals were recorded at 510 nm following excitation at 485 nm. The calcein release percentage at various AapA1 concentrations was calculated as follows: Calcein release percentage = (experimental group – negative control) / (positive control – negative control) × 100%

#### Pore blocking experiments

The pore blocking experiments were performed as described previously with slight modifications ^42^. Briefly, 40 mM polyethylene glycol (PEG) of different molecular weights (400, 600, 1000, 2000, and 4000 Da) was dissolved in PBS; 0.5% DMSO without PEG and 0.5% Triton X-100 without PEG were used as negative and positive controls, respectively. Liposomes loaded with calcein were diluted in PBS containing different PEG fractions, mixed with an equal volume of 80 μM AapA1, and incubated at 37 °C for 10 min in the dark. Calcein fluorescence signals were recorded following the method described in the subsection “Calcein release assay.”

#### Vitamin D3 release assay

Vitamin D3-enriched liposomes were mixed with 40 μM peptides and incubated for 10 min at 37 °C; 0.5% DMSO was used as a negative control. Liposomes were removed from the samples by centrifugation using a 100-kDa ultrafiltration tube, and the filtrates were collected. Vitamin D3 content in the samples was quantified by spectrophotometry at 254 nm. The absorbance value of the standard sample was used for constructing the standard curve. The sample concentration was calculated from the standard curve.

#### Statistical analysis

Data are expressed as mean (n ≥ 3) ± standard deviation (SD) and are shown from single representative experiments repeated independently at least two times for each condition. ANOVA was used for comparing more than two independent groups. Differences with P < 0.05 were considered statistically significant. All statistical analyses were conducted using GraphPad Prism 8.0 software.

### 2. Classical molecular dynamics (CMD) simulation setup for multimeric AapA1-membrane systems

AapA1 protein sequence (MATKHGKNSWKTLYLKISFLGCKVVVLLKR) and secondary structure characteristics (PDB ID: 6GIG) ^10^ are shown in Figures 1A and 1B. The S9-L28 residues of the AapA1 monomer protein formed a stable transmembrane helix ^10^. The multimeric AapA1 protein structures of the tetramer, pentamer, hexamer, heptamer, octamer, and nonamer were predicted by ADMFold-Multimer software (https://zhanggroup.org/DMFold/) ^43^, and the top 5 structures are shown in Figure S1. The top 1 model with the highest predicted TM score (pTM-score) for the multimer was selected as the initial configurations (shown in Figures 1D) for these simulations. These multimeric AapA1 toxin proteins were tightly packed in a parallel multimeric helix bundle (N terminus in the upper leaflet; shown in Figures 1D), except for the antiparallel tetramer. All these conformations showed a common feature: aggregation of the K16 and K23 residue side chains opposite to each other and facing the channel pore. The POPE: POPG ratio in the membrane bilayer was 3:1, with a total of 144 POPE and 48 POPG molecules per simulation system. This membrane bilayer is a commonly used computer model for the inner membrane of gram-negative bacteria. Multimeric AapA1 insertion into the membrane bilayer was accomplished using the CHARMM-GUI web server (http://charmm-gui.org/) ^44, 45^. CHARMM36 force fields ^46, 47^ were used for AapA1 and the POPE: POPG 3:1 bilayer, and the TIP3P model was used for water molecules, with neutralization by 0.15 M NaCl. For each system, the simulation box was approximately 8.3 × 8.3 × 12 nm^3^ in size and contained approximately 85,000 atoms. First, steepest descent energy minimization was performed to relax the simulation system. Subsequently, 1 ns NVT and NPT simulations were conducted to relax the solvent molecules and lipid bilayer. All simulations were performed with the Nose−Hoover algorithm ^48^ at 310 K temperature and 1 bar pressure by using the semi-isotropic Parrinello−Rahman method ^49^. The Particle-mesh Ewald method ^50^ was used to calculate the long-range electrostatic interactions with a grid spacing of 1.2 nm. The Lennard−Jones interaction was calculated using the force-based switching function with a cutoff of 1 nm. The bond length was constrained using the LINCS algorithm ^51^. Three sets of CMD simulations were performed for each multimeric AapA1-membrane system (simulation details shown in Table 1).

**Figure 1.**
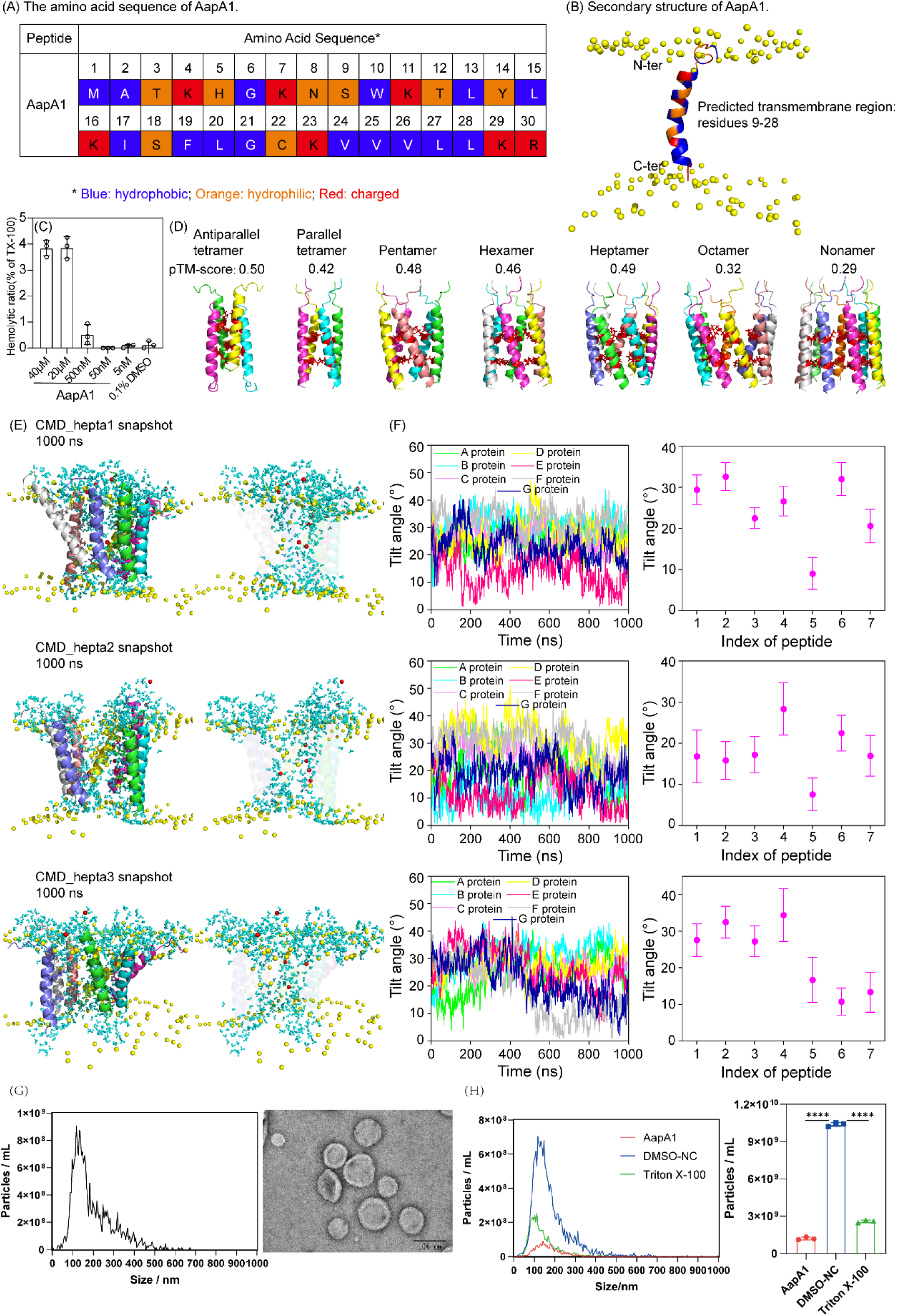
Structure of AapA1 toxin protein. (A) The amino acid sequence of AapA is presented, with different residue types represented by distinct colors: blue for hydrophobic residues, orange for hydrophilic residues, and red for charged residues. (B) The transmembrane structure of AapA1 is illustrated, highlighting its amphipathic α-helical configuration. The phosphorus atoms are shown in yellow spheres and the corresponding amino acid residues are color-coded according to the scheme used in (A). (C) Haemolytic activity of different concentrations of AapA1 for erythrocytes. (D) The multimeric AapA1 toxin proteins structures and corresponding pTM-score predicted by software ADMFold-Multimer. The K-16 and K-23 residues are shown in red sticks. (E) The 1000th ns snapshots of the Heptamer AapA1 and POPE/POPG (3:1) structure of CMD_hepta simulations. In the snapshots, the following colors are used for the seven AapA1 proteins: A protein, green; B protein, cyan; C protein, lightmagenta; D protein, yellow; E protein, salmon; F protein, gray90; G protein, slate; the lipid phosphorus atoms are shown as yellow spheres, water are shown as cyan sticks, and chloridions are shown as red spheres. (F) The time series of fluctuation of tilt angles for each AapA1 protein (left) and the average tilt angles for each AapA1 protein during the last 200 ns (right). (G) Nanoparticle tracking analysis (NTA) of 3POPE:1POPG liposomes (left) and images under electron microscope (right), scale bar, 100 nm. (H) 40 μM AapA1 were incubated with 3POPE:1POPG liposomes for 10 min at 37°C, 0.5% DMSO as a negative control, and 0.5% TritonX-100 as a positive control. The left panel shows the results of NTA assay and the right panel shows the sum of 100-200 nm liposome concentrations. The experiments were performed in triplicate, and representative data are shown as mean ± SD. Statistical analyses were performed using unpaired *t*-test with GraphPad Prism 8.0 software, \*\*\*\**P* < 0.0001.

**Table 1.**
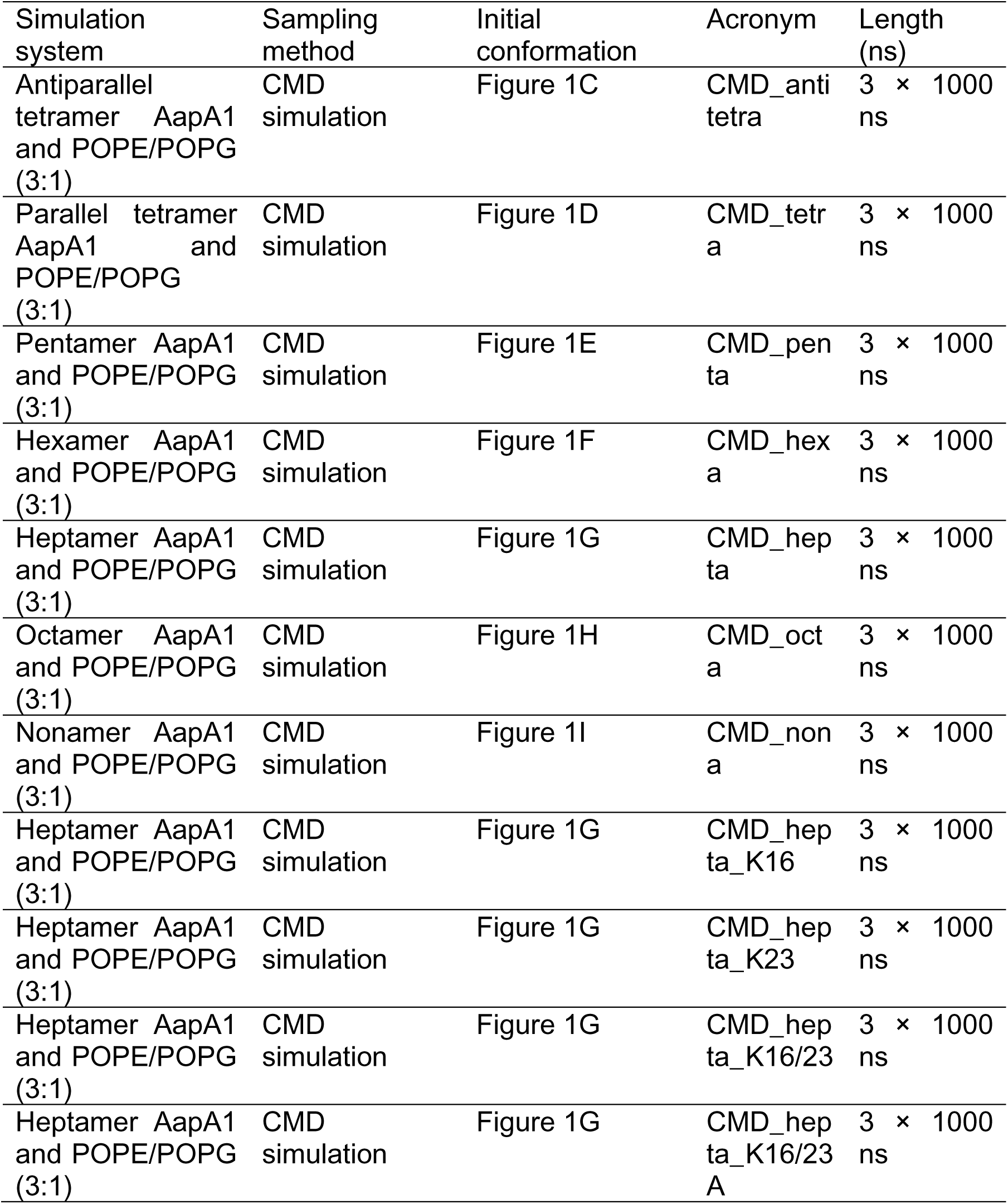
Summary of the details of multimeric AapA1 toxin proteins-membrane simulations.

| Simulation system | Sampling method | Initial conformation | Acronym | Length (ns) |
| --- | --- | --- | --- | --- |
| Antiparallel tetramer AapA1 and POPE/POPG (3:1) | CMD simulation | Figure 1C | CMD_anti tetra | 3 × 1000 ns |
| Parallel tetramer AapA1 and POPE/POPG (3:1) | CMD simulation | Figure 1D | CMD_tetra | 3 × 1000 ns |
| Pentamer AapA1 and POPE/POPG (3:1) | CMD simulation | Figure 1E | CMD_penta | 3 × 1000 ns |
| Hexamer AapA1 and POPE/POPG (3:1) | CMD simulation | Figure 1F | CMD_hexa | 3 × 1000 ns |
| Heptamer AapA1 and POPE/POPG (3:1) | CMD simulation | Figure 1G | CMD_hepta | 3 × 1000 ns |
| Octamer AapA1 and POPE/POPG (3:1) | CMD simulation | Figure 1H | CMD_octa | 3 × 1000 ns |
| Nonamer AapA1 and POPE/POPG (3:1) | CMD simulation | Figure 1I | CMD_nona | 3 × 1000 ns |
| Heptamer AapA1 and POPE/POPG (3:1) | CMD simulation | Figure 1G | CMD_hepta_K16 | 3 × 1000 ns |
| Heptamer AapA1 and POPE/POPG (3:1) | CMD simulation | Figure 1G | CMD_hepta_K23 | 3 × 1000 ns |
| Heptamer AapA1 and POPE/POPG (3:1) | CMD simulation | Figure 1G | CMD_hepta_K16/23 | 3 × 1000 ns |
| Heptamer AapA1 and POPE/POPG (3:1) | CMD simulation | Figure 1G | CMD_hepta_K16/23 A | 3 × 1000 ns |

### 3. WT-BEMetaD and CMD simulations for monomer AapA1-membrane systems

We performed a 2 µs WT-BEMetaD simulation for the membrane systems containing monomeric AapA1 and Figure 1B shows the initial conformation. The simulation box for monomer AapA1-POPE: POPG [3:1] was approximately 6.3 × 6.3 × 14 nm^3^ in size, with a total of 96 POPE and 32 POPG molecules per simulation system. The remaining simulation parameters were the same as mentioned above. The WT-BEMetaD simulation ^52^ was performed using the GROMACS 2018 software package ^53^ patched with the PLUMED 2.5 plugin ^54^. To analyze the localization of K16 and K23 in the membrane, we used the total number of contacts (collective variable [CV] was defined as the coordination number in PLUMED) between the K16/K23 residues and the phosphate group atoms in either the top or bottom leaflet of the membrane as the two (CVs). Table 2 shows the specific description of CVs. The well-tempered approach was utilized in all WT-BEMetaD simulations, with the addition of the bias potential every 10 ps. The initial Gaussian height and the width for the contact number were set to 1 kJ/mol and 10, respectively. A bias factor of 10 was utilized. Additionally, three sets of 1000 ns CMD simulations were initiated from the local energy minimum states obtained from the WT-BEMetaD simulation, as shown in Table 2. The simulation parameters were the same as mentioned previously. All simulated systems were equilibrated for 800 ns, and subsequent 200 ns simulations were used for the analysis.

**Table 2.** Summary of the details of single AapA1 toxin protein-membrane simulations.

| Simulation system | Sampling method | Initial conformation | Collective variables (CVs) | Acronym | Length (ns) |
| --- | --- | --- | --- | --- | --- |
| 1 AapA1 and POPE/POPG (3:1) | WT-BEMetaD simulation | Figure 1B | CV1: Contact number between K16/K23 and the top phosphate group.<br>CV2: Contact number between K16/K23 and the bottom phosphate group. | WT1 | 2 replica x 2000 ns |
| 1 AapA1 and POPE/POPG (3:1) | CMD simulation | representative conformation of region A in Figure 7 |  | CMD_SP1 | 3 x 1000 ns |
|  |  | representative conformation of region B in Figure 7 |  | CMD_SP2 | 3 x 1000 ns |
|  |  | representative conformation of region C in Figure 7 |  | CMD_SP3 | 3 x 1000 ns |

## Results

### 1. Multimeric AapA1 can stably exist in the membrane and form pores

The AapA1 peptide was synthesized in solid phase from the amino acid sequence and the predicted transmembrane region of the secondary structure is residues 9-28 (Figure 1A-B). AapA1 exhibited almost no hemolytic effect on erythrocytes, and AapA1 at a very high concentration (40 μM) caused <4% hemolytic damage to erythrocytes (Figure 1C). This result suggests that AapA1 might cause minimal damage to mammalian cells. CMD simulations were then conducted to investigate the structural characteristics of seven different multimeric AapA1-membrane systems (antiparallel-tetramer, tetramer, pentamer, hexamer, heptamer, octamer, and nonamer) (Figure 1D and S1). We discuss the heptameric AapA1-membrane system as an example here. The 1000th ns snapshots from the CMD_hepta simulations of the AapA1 heptamer in POPE: POPG (3:1) liposomes and tilt angles for each protein (shown in Figure 1E-F). The state of each protein was defined according to previous studies ^55^, and tilt angles were categorized as follows: 60°– 120° as surface state (S-state), 30°–60°/120°–150° as tilted state (T-state), and 0°–30°/150°–180° as inserted state (I-state). In the initial conformation, the tilt angle of the heptameric AapA1 protein was approximately 24°. During the simulations, the range of tilt angles for the seven proteins was 0–40°, and the proteins mainly existed in the I-state. The minimum and maximum tilt angles of AapA1 were approximately 8° and 35°, respectively. The microsecond-long MD simulations revealed the existence of relatively large-amplitude motions of the transmembrane helices of the AapA1 protein within the bilayer membrane.

Liposomes of approximately 120 nm in size were prepared from bacterial cell membrane components (POPE: POPG = 3:1) by using the lipid extrusion method (Figure 1G). AapA1-induced disruption of the intact vesicle structure of liposomes was detected by NTA. The degree of disruption of liposome integrity by 40 μM AapA1 was approximately 87% (Figure 1H). The abovementioned experimental results confirmed that AapA1 forms pores on the POPE: POPG (3:1) liposomes. However, it remains unclear how AapA1 affects cell membrane integrity and mediates toxicity.

Figures 2 show the dynamic changes of the heptameric AapA1-membrane system. As observed in the three snapshots of CMD_hepta simulations, some phospholipid head groups in the upper and lower lipid layers moved to the center of the membrane. The average distances of the individual phosphorus atoms from the center of membrane (COM) of the lipid bilayer were calculated from the last 200 ns trajectories in the CMD_hepta simulations (Figures 2A-C). The bilayer membrane showed relatively visible deformations, with approximately 4–8 phospholipid head groups embedded in the hydrophobic lipid core. The membrane leakage effect induced by heptamerized AapA1 proteins was measured by calculating the Z-dependent chloride ions (Figure 2D), water mass density (Figure 2E), and the average number of water molecules in the different regions of the lipid bilayer across 200 ns conformations. The membrane was centered at Z = 0, and the bilayer core, represented by a 0.9-nm region in the mid-plane of the bilayer, contained approximately 60 water molecules and 2–4 chloride ions at the mass density of 30 kg/m^3^ and 2–3 kg/m^3^, respectively, in the CMD_hepta simulations (Table 3). The Deuterium order parameters (|S_CD_|) of acyl chains was calculated using the Membrainy tool ^56^ for the pure POPE: POPG (3:1) lipid bilayer and CMD_hepta simulations (Figures 2F-G). Acyl chains were more disordered in the CMD_hepta simulations than in the pure POPE: POPG membrane. Hence, the heptamerized AapA1 protein with the transmembrane structure can exist stably in the membrane, resulting in bilayer membrane deformation and water/chloride ion pore formation. Calcein release assay was conducted to determine the concentration-dependent effects of AapA1 on cell membrane permeability (Figure 2H). The Einstein–Stokes radius (ESR) for PEG is closely correlated with its molecular weight; the higher the molecular weight, the larger is the molecular ESR^57^. We hypothesized that the diameter of pores formed by AapA1 on liposomes was 2–3 nm (Figure 2I). All of the above evidence supports that AapA1 affects membrane permeability through pores in the membrane.

**Figure 2.**
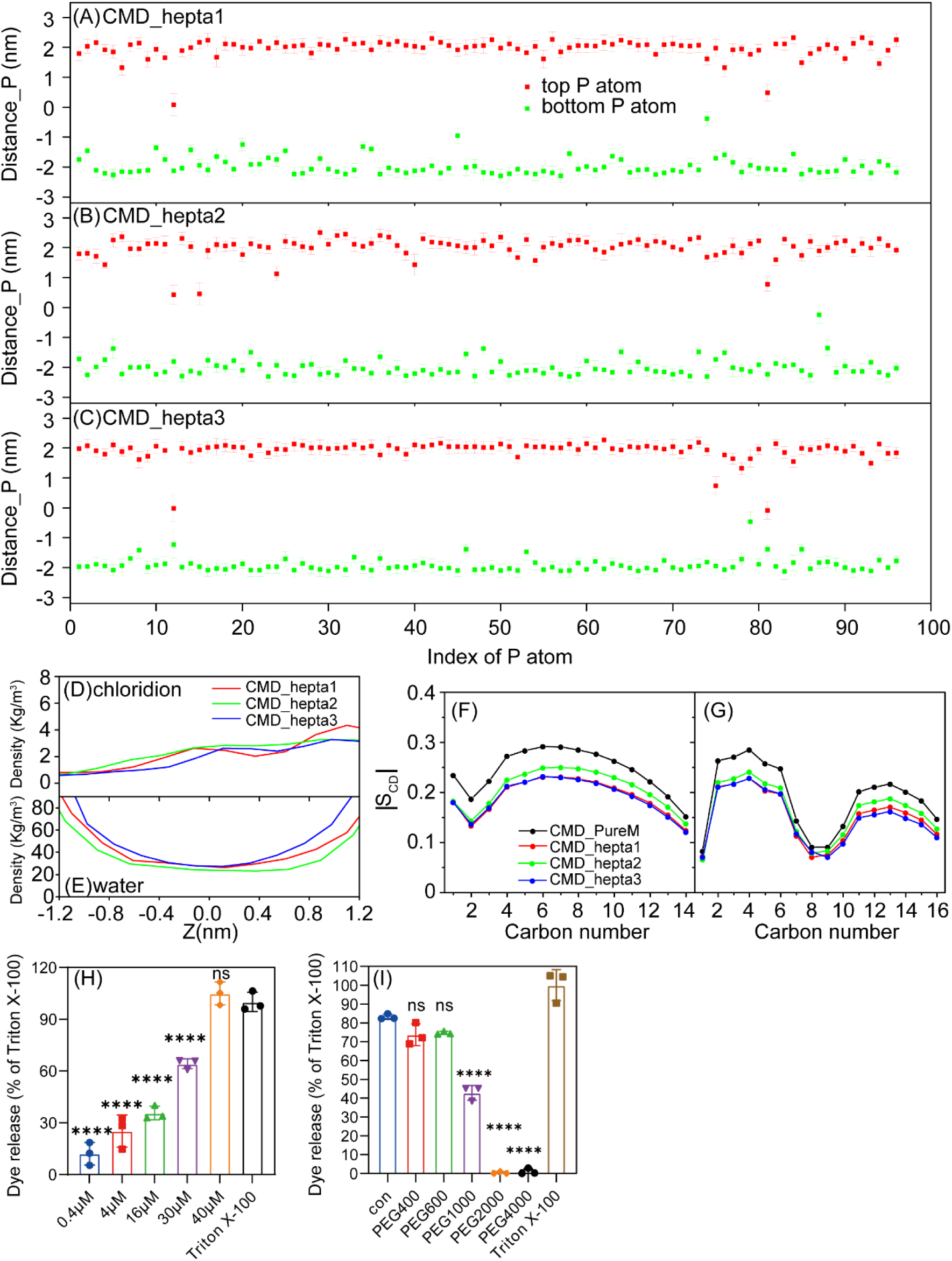
(A-C) The average distance of each phosphorus atom from the COM of the lipid bilayer during the last 200 ns of the trajectories in the CMD_hepta simulations. “Distance_P” represents the average distance for phosphorus atoms. (D-E) Mass density of chloridions and water in the CMD_hepta simulations. Deuterium order parameters (|SCD|) of the saturated (F) and unsaturated (G) lipid acyl chains in the CMD_hepta simulations. (H) Different concentrations of AapA1 were incubated with calcein-containing liposomes at 37°C for 10 min, with 0.5% DMSO as negative control and 0.5% TritonX-100 as positive control. The calcein fluorescence signal was recorded at 510 nm upon excitation at 485 nm. (I) Pore blocking assay with different sized PEG particles. 40 mM of different sized PEG particles were incubated with fluorescent liposomes mixed with a final concentration of 40 μM of AapA1, incubated at 37°C for 10 min. The calcein fluorescence signal was recorded at 510 nm upon excitation at 485 nm. The experiments were performed in triplicate, and representative data are shown as mean ± SD. Statistical analyses were performed using unpaired *t*-test with GraphPad Prism 8.0 software, ns, *P* ≥ 0.05, \*\*\*\**P* < 0.0001.

**Table 3.**
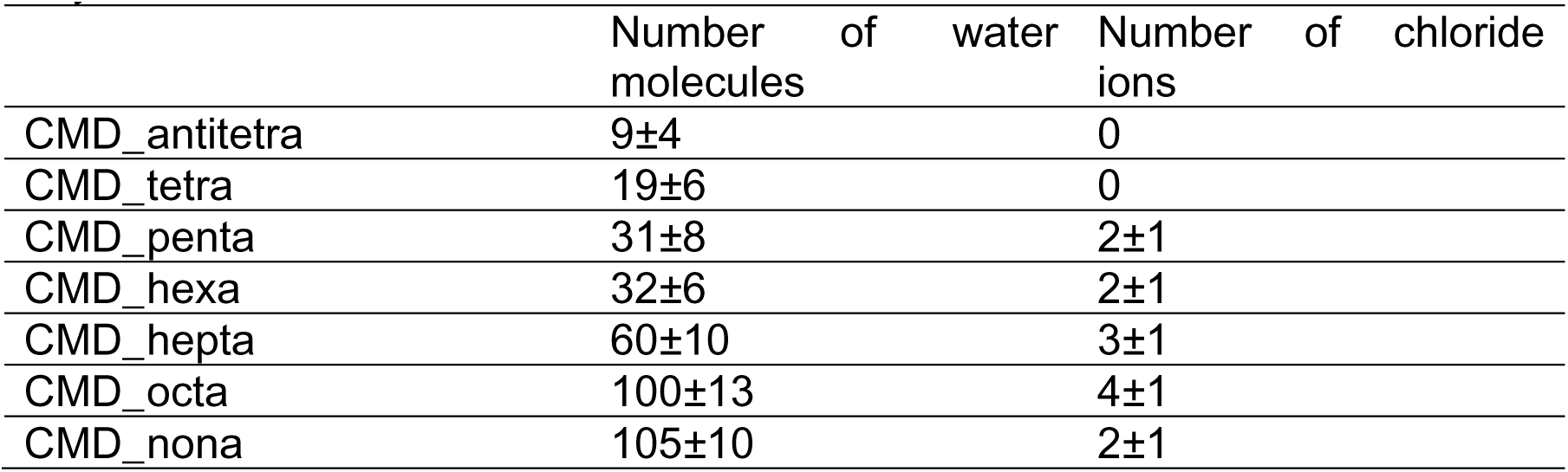
The number of water molecules and chloride ions in the mid-plane of the bilayer.

The AapA1 protein contains only positively charged/hydrophobic residues and no negatively charged residues. The electrostatic interaction between AapA1 and the membrane mainly involves the positively charged residues K4, K7, K11, K16, K23, K29, and R30. K4 and K7 residues are mainly found on the membrane surface. The average distance of each COM of the K11, K16, K23, K29, and R30 residue side chains from the COM of the lipid bilayer was calculated during the last 200 ns of the trajectories in the CMD_hepta simulations (Figure 3A-C). The K11 residue always interacted with the upper membrane, and the K29 and R30 residues always interacted with the lower membrane; however, the K16 and K23 residues did not interact with a particular layer of the lipid bilayer. The distance between the K16 or K23 residue and the center of the membrane fluctuated over a wide range from –1.0 to 1.6 nm (Figure 3A-C).

**Figure 3.**
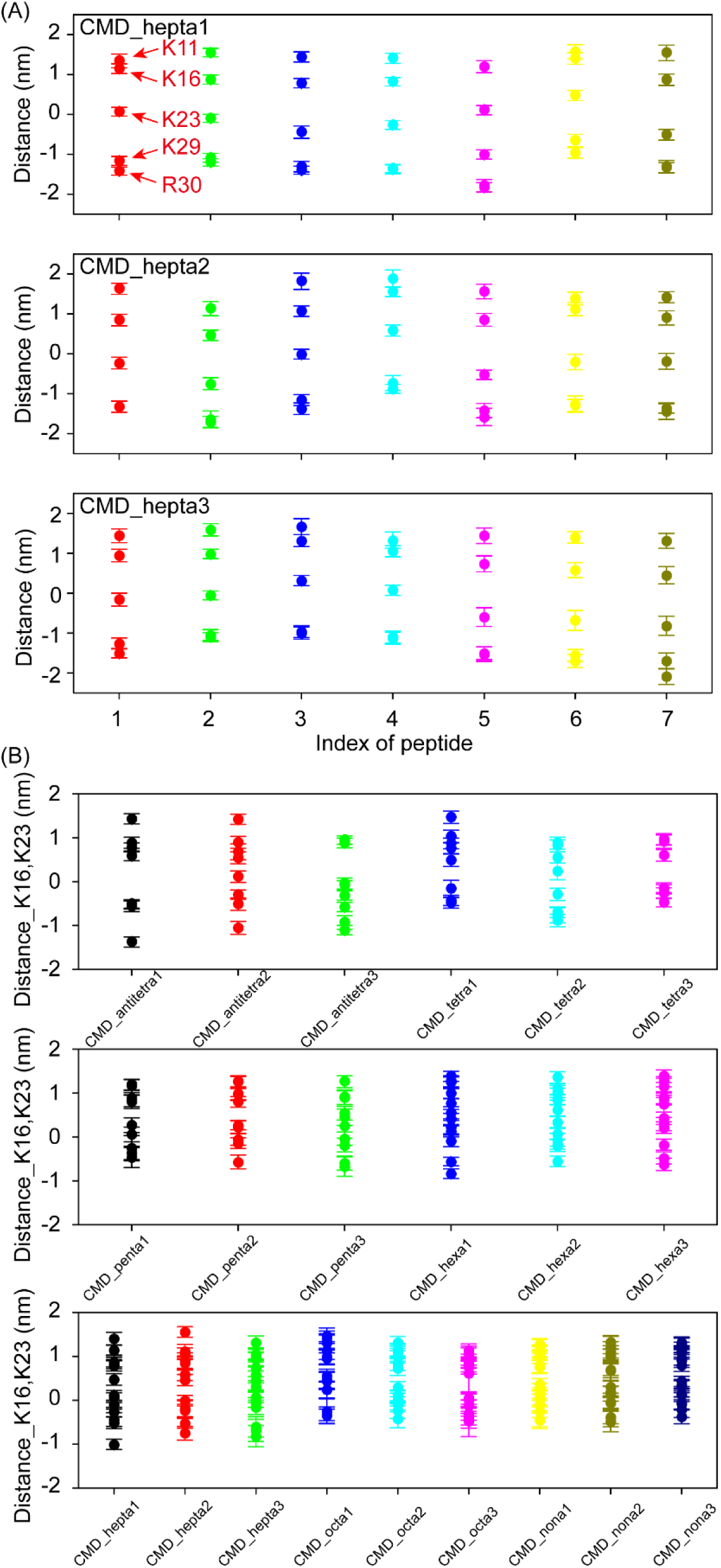
(A) The average distance of each COM of the K11, K16, K23, K29 and K30 residue side chain from the COM of the lipid bilayer during the last 200 ns of the trajectories in the CMD_hepta simulations. (B) The average distance of each COM of the K16 and K23 residue side chain from the COM of the lipid bilayer (labeled as Distance_K16, K23) during the last 200 ns of the trajectories in 21 multimeric AapA1 proteins simulations.

The structural characteristics and dynamic changes in the lipid bilayer for the other six multimeric AapA1 proteins are shown in Figures S2–S7: antiparallel-tetramer (Figures S2), tetramer (Figures S3), pentamer (Figures S4), hexamer (Figures S5), octamer (Figures S6), and nonamer (Figures S7). These multimeric AapA1 protein simulations showed three significant findings as follows. Firstly, AapA1 existed stably as antiparallel-tetramer, tetramer, pentamer, hexamer, heptamer, octamer, and nonamer. The multimeric AapA1-lipid membrane interaction was not only strongly associated with membrane deformation but also limited to pore-induced membrane permeabilization. Secondly, with the increase in the number of multimeric AapA1 proteins (from 4 to 9), |S_CD_| decreased and the Z-dependent water mass density (Figure S8) and the number of water molecules in the mid-plane of the bilayer (Table 3) gradually increased; the mass density of Z-dependent chloride ions and the number of chloride ions in the mid-plane of the bilayer increased and were balanced to a certain extent. Thirdly, the distance between the K16/K23 residues and the center of the membrane showed a wide range of fluctuation in all multimeric AapA1 protein-membrane simulations (Figure 3D-F). The side chains of K16/K23 residues were arranged as a ladder in the cell membrane. AapA1 proteins self-assembled into water-soluble α-helical barrels in the membrane, wherein by lining the interior with positive residues K16 and K23 and the exterior with hydrophobic residues, respectively. These barriers were restructured to form a polar pore. Hence, the determination of the preferred conformational state of AapA1 in the POPE: POPG membrane is essential for further analysis and for understanding its mechanism of action and functional implications.

### 2. Swing motions of the K16/K23 side chains in the transmembrane helix are critical to water permeability

The microsecond-long CMD simulations of the multimeric AapA1-membrane systems revealed two interesting findings: (1) relatively large fluctuation in the orientation of transmembrane helices and (2) relatively large variation in the distance between K16/K23 residues and the center of the membrane. To investigate the relationship between the bobbing of the K16/K23 side chains and the orientation of transmembrane helices and how this relationship affects water permeation, we designed three different constrained simulation systems (details are provided in Table 1) and compared them with the CMD_hepta simulations (all atoms could move freely in the systems). Position restraints were added on the side chain atoms of K16, K23, and K16/K23 in the CMD_hepta_K16 simulations, CMD_hepta_K23 simulations, and CMD_hepta_K16/K23 simulations, respectively. Figure 4A-C shows the 1000th ns snapshots of the three restrained simulations and tilt angles for each AapA1 protein. In these three constrained simulations, the approximate average tilt angles were 15°, 24°, and 19°, respectively (Figure 4E-G). In the CMD_hepta_K23 simulations, two proteins had a larger tilt angle of approximately 30°. In the CMD_hepta_K16/K23 simulations, the range of fluctuations in the orientation angle of transmembrane helices was smaller than that observed in the CMD_hepta_K16 and CMD_hepta_K23 simulations. The microsecond-long restrained MD simulations revealed that the fluctuation range of the tilt angle was reduced because of the constrained side chain of K16/K23, particularly K16. We also investigated the effect of K16/K23A mutation on the structural characteristics of heptamer in the CMD_hepta_K16/K23A simulations (Figure 4D and 4H). Figures S9 show structural characteristics and dynamic changes in the lipid bilayer for heptamer AapA1 with K16/K23A mutation. The results of the CMD_hepta_K16/K23A simulations were completely different from those of the CMD_hepta simulations. The center of the membrane had no phospholipid head group, and its water density was close to zero.

**Figure 4.**
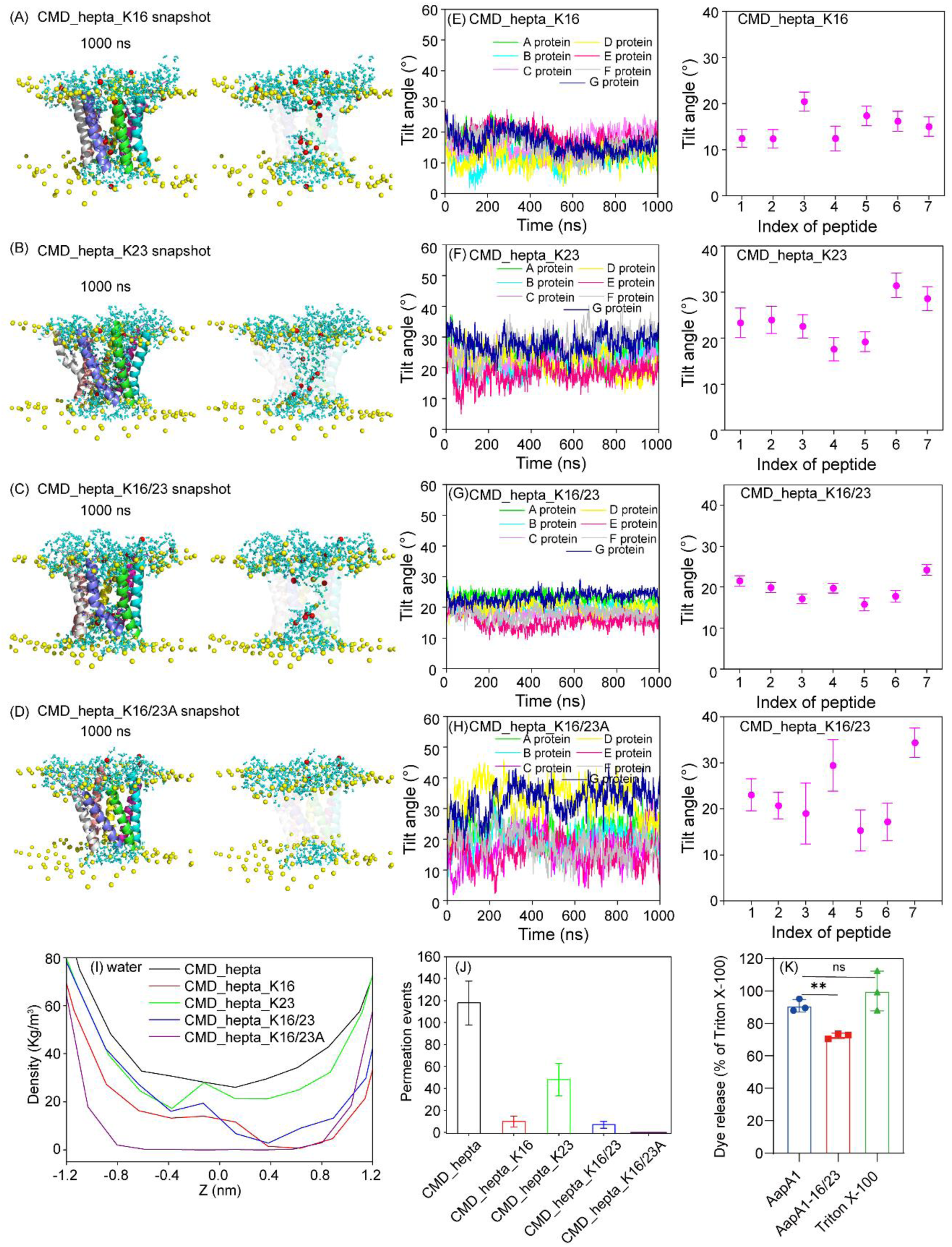
(A-D) The 1000th ns snapshots of the Heptamer AapA1 and POPE/POPG (3:1) structure of CMD_hepta_K16, CMD_hepta_K23 CMD_hepta_K16/23 and CMD_hepta_K16/23A simulations. (E-H) The time series of fluctuation of tilt angles for each AapA1 protein (left) and the average tilt angles for each AapA1 protein during the last 200 ns (right). The colors are the same as in Figure 1E-F. (I) The Z-dependent water mass density and (J) the average amount of permeated water molecules passing the pore per 100 ns. (K) Final concentrations of 40 μM AapA1 or AapA1-16/23 mutant proteins, respectively, were incubated with calcein-containing liposomes at 37°C for 10 min, with 0.5% DMSO as a negative control and 0.5% TritonX-100 as a positive control. The calcein fluorescence signal was recorded at 510 nm upon excitation at 485 nm. The experiments were performed in triplicate, and representative data are shown as mean ± SD. Statistical analyses were performed using unpaired *t*-test with GraphPad Prism 8.0 software, ns, *P* ≥ 0.05, \*\**P* < 0.01.

The Z-dependent water mass density (Figure 4I) and the amount of permeated water molecules passing through the pore (Figure 4J) in the different simulation systems were calculated. Water permeation was defined as the complete crossing of a water molecule from one side of the lipid bilayer to the other side. The free dynamic CMD_hepta simulation systems exhibited the highest water translocation efficiency. An average of approximately 118 water molecules were translocated in both directions every 100 ns. A comparison of translocated water molecules revealed that only approximately 10, 48, 7, and 0 water molecules were translocated in the CMD_hepta_K16, CMD_hepta_K23, CMD_hepta_K16/K23, and CMD_hepta_K16/K23A simulations, respectively. Mutations in amino acids at K16 and K23 (AapA1-16/23) reduced the membrane permeability function of AapA1 (Figure 4K). The comparison of water translocation efficiencies of these five dynamics systems indicates that the swing motion of the K16/K23 side chains in the transmembrane helix is critical for water transfer.

### 3. Three possible conformational states of the single AapA1 protein-membrane complex because of the different orientations of the K16/K23 side chains

The abovementioned CMD simulations provide a meaningful and realistic representation of the structural characteristics of multimeric AapA1 proteins in the cell membrane; however, these investigations were fundamentally limited by the sampling efficiency of CMD simulations. Hence, WT-BEMetaD simulation was conducted to investigate the effect of K16/K23 side chain motions on the structure of AapA1 in the cell membrane. The possible multiple conformational states of the single AapA1 protein in the membrane were analyzed from the two-dimensional (2D) free energy landscapes (FELs) generated using the two CVs (Figure 5A) by reweighting WT1 simulation using the sum_hills tool ^58^. Figure 5A depict the 2D FELs as a function of the number of contacts among residues K16/K23 and the atoms of the phosphate groups in the upper or lower leaflet of the membrane in WT1 simulation. Three local energy minima are shown in Figure 5A: region A (CV1 = 60, CV2 = 170) with a free energy of 0 kJ/mol; region B (CV1 = 110, CV2 = 100) with a free energy of 5.6 kJ/mol, and region C (CV1 = 140, CV2 = 60) with a free energy of 15.0 kJ/mol. The AapA1 protein was inserted into the lipid bilayer membrane, and the protein aligned itself along the Z-axis of the bilayer. The three representative conformations mainly differed in the orientation of the K16 and K23 side chains. In region A, the K16 and K23 side chain residues were oriented toward the bottom layer of the membrane. In region B, the K16 side chain residues were oriented toward the upper layer of the membrane, while the K23 side chain residues were oriented toward the lower layer of the membrane. In region C, the K16 and K23 side chain residues were oriented toward the upper layer of the membrane. The K16 and K23 side chain residues can orient toward the upper or lower layer of the membrane. Although the K16 and K23 side chain residues are positively charged, they can still exhibit bobbing motion inside the bilayer membrane.

**Figure 5.**
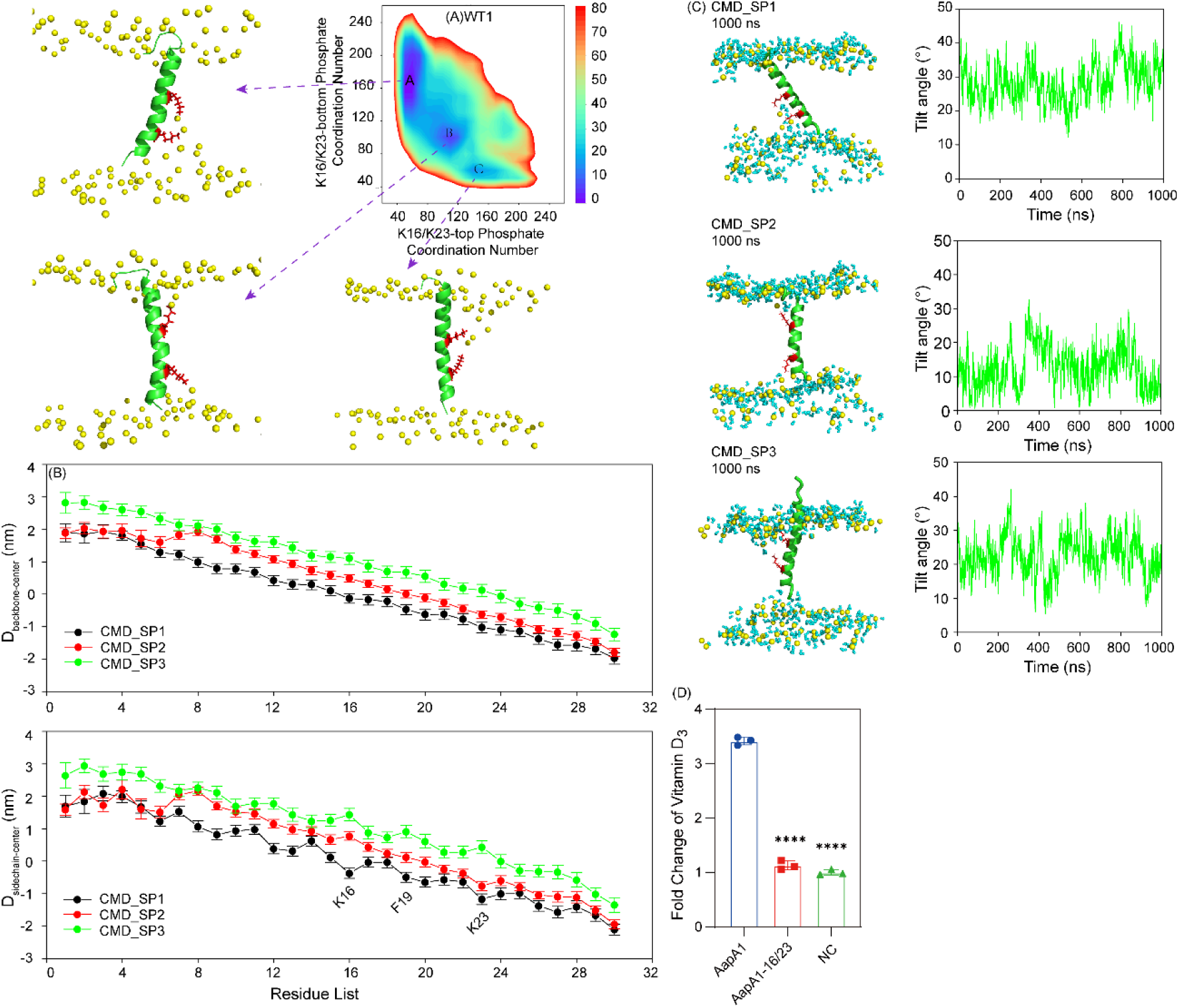
(A) Two-dimensional free energy landscapes depicting the relationship between CV1 and CV2 for WT1 simulation. The labeled local minima and the representative conformation are denoted as A, B, and C. K16 and K23 residues are shown in red stick. The global minimum is assigned a free energy value of zero. (B) The average distance and variance (up) of each residue backbone to the membrane center and (down) of each residue side chain to the membrane center. (C) The 1000th ns snapshots of the single AapA1-membrane structure of CMD_SP1, CMD_SP2 and CMD_SP3 simulations (left) and the time series of fluctuation of tilt angles for AapA1 protein (right). (D) Final concentrations of 40 μM AapA1 or AapA1-16/23 mutant proteins, respectively, were incubated with vitamin D3-containing liposomes at 37°C for 10 min, with 0.5% DMSO as a negative control. samples were centrifuged in 100K ultrafiltration tubes, and the filtrate was assayed for VD3 at 254 nm. The experiments were performed in triplicate, and representative data are shown as mean ± SD. Statistical analyses were performed using unpaired *t*-test with GraphPad Prism 8.0 software, \*\*\*\**P* < 0.0001.

The abovementioned three local minima (A, B, and C) and their representative conformations (Figure 5A) served as the initial conformations for the CMD_SP simulations (details are presented in Table 2). To compare the structural characteristics of the AapA1 protein in these three distinct conformational states, we calculated the position of each residue in the three sets of CMD_SP simulations. The distance between the backbone or side chain of each residue and the center of the membrane was calculated (Figure 5B). These three conformational states primarily differed in the orientation of the K16 and K23 side chains, either toward the upper or lower cell membrane. In the CMD_SP1, CMD_SP2, and CMD_SP3 simulations, the K16 and K23 side chains were located at approximately –0.4 nm and –1.2 nm, 0.8 nm and –0.8 nm, and 1.4 nm and 0.4 nm, respectively, from the center of the membrane. The three different transmembrane structures of AapA1 in CMD_SP1, CMD_SP2, and CMD_SP3 simulations remained highly stable throughout the 1000 ns trajectory. The snapshots of the AapA1-membrane complex at the 1000th ns in the CMD_SP1, CMD_SP2, and CMD_SP3 simulation trajectories were shown (Figure 5C).

The binding free energies between AapA1 and the POPE: POPG (3:1) bilayer were evaluated for the last 200 ns of CMD_SP simulations (Table 4) by using the g_mmpbsa software ^59^. Table 4 shows the total binding free energy involving residues 1–30 or residues 9–30 and the membrane. Regardless of whether K16 or K23 interacted with the upper or lower lipid membrane, slight differences were noted in the electrostatic energy and polar solvation energy, with no significant differences in van der Waals energy and nonpolar solvation energy. The total binding free energy of the AapA1-POPE: POPG (3:1) lipid bilayer in CMD_SP1 simulations was slightly lower than that in CMD_SP2 and CMD_SP3 simulations. Vitamin D3 showed a desirable anti-*H. pylori* effect in both *in vitro* and *in vivo* experiments, even against antibiotic-resistant strains ^9^. Our experiments demonstrated that the pores formed by AapA1 on the membrane allowed the permeation of vitamin D3, and this membrane permeability was controlled by amino acids at K16 and K23 (Figure 5D). We hypothesized that this function possibly facilitates exocytosis of intracellular vitamin D3 in *H. pylori*, thereby increasing its resistance to antibiotics.

**Table 4.**
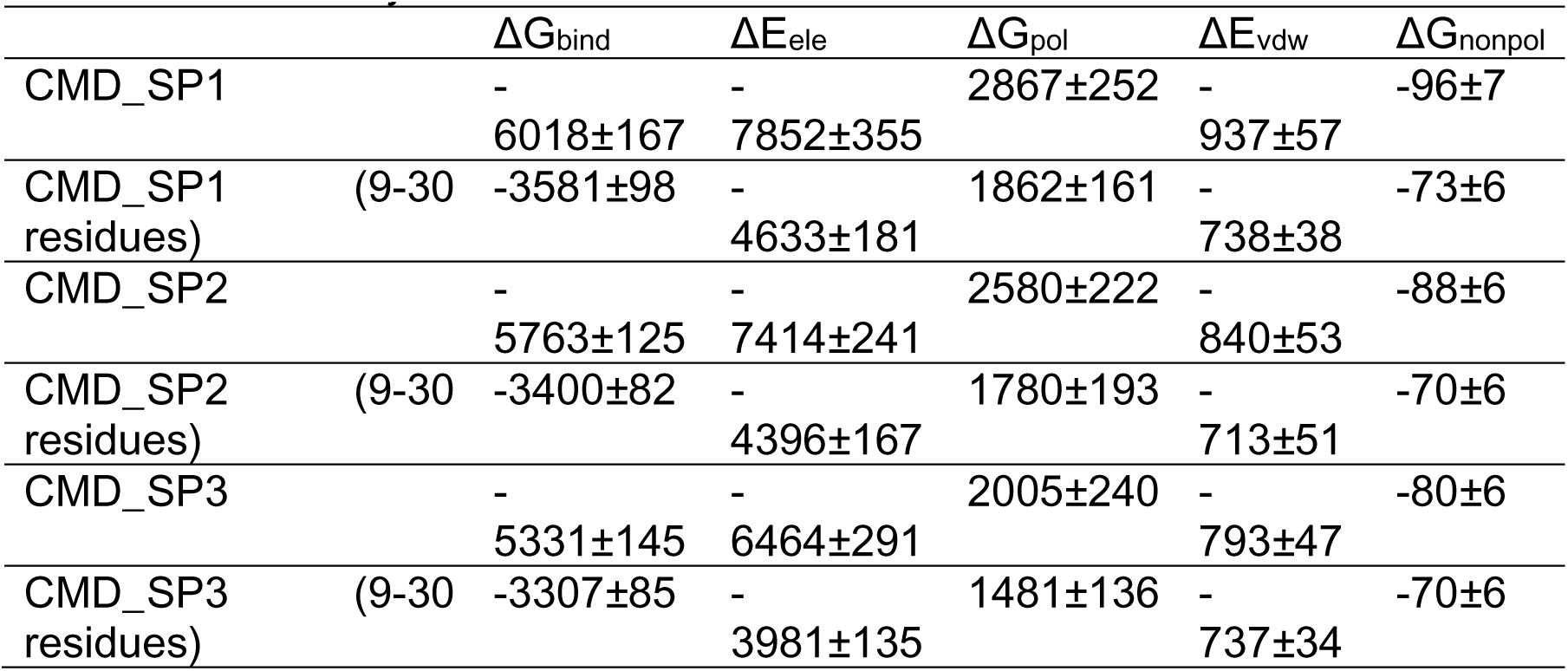
Binding free energy (kJ/mol) between the monomeric AapA1 toxin protein and the bilayer membrane in the CMD_SP1-3 simulations during the last 100 ns of MD simulation trajectories.

## Discussion

The type I TA system in bacteria has crucial biological functions related to both growth regulation and antibiotic resistance. Previous studies have confirmed that AapA1 expression in *H. pylori* induces a rapid transition from the helical form to the coccoid form, leading to growth suppression and entry into an active dormant state, which is more prolonged than the retention state. AapA1 is the first effector molecule identified to promote the formation of coccoid bacterial forms by targeting the intracellular membrane of *H. pylori*. It acts against the intracellular membrane of *H. pylori*, without causing membrane disruption.

In the present study, *in vitro* experiments were combined with MD simulations to analyze the structural characteristics and mechanism of action of AapA1 on the cell membrane. AapA1 induced minimal damage to mammalian erythrocytes, and a very high concentration (40 μM) of AapA1 caused <4% hemolytic damage to erythrocytes (Figure 1C); this finding suggests that AapA1 exhibits selective transmembrane activity on the target bacteria cell membrane. Residues S9-L28 of monomeric AapA1 have a stable transmembrane helical structure [4]. We further demonstrated that the S9-L28 residues of AapA1 exist as tightly packed multimeric helical bundles, which allows the K16 and K23 side chains to orient toward the inner side of the channel (Figures 1D). We investigated different multimeric AapA1 protein-membrane systems by using a series of CMD simulations and found that multimeric AapA1 can stably exist in the simulated bacterial cell membrane system; however, the transmembrane helical region had large-amplitude motions within the bilayer, which caused membrane collapse (Figure 1E-F). This finding suggests that the transmembrane helical region of multimeric AapA1 is strongly perturbative for the bilayer membrane. A similar phenomenon was observed in *in vitro* experiments, wherein AapA1 showed active disruption of liposomes mimicking bacterial cell membrane components (Figure 1H).

CMD simulations were conducted to determine the physical properties of the transmembrane helical region of multimeric AapA1; the results showed that the pores formed by the transmembrane helical region in the lipid bilayer allowed the passage of water (Figure 2A-G). We then constructed a liposome-hydrophilic fluorescent dye model and found that AapA1 induced the release of hydrophilic dyes from liposomes and showed a concentration-dependent effect (Figure 2H). These results clearly indicate that multimeric AapA1 is involved in the formation of hydrophilic pores on the bacterial cell membrane. Taken together with the simulation results, we hypothesized that AapA1 can exist stably in the cell membrane as α-helices through multimers and form hydrophilic pores of different sizes. Pore blocking experiments with PEG of various molecular sizes showed that multimeric AapA1-formed pores were approximately 2 nm in size (Figure 2I). This finding suggests that multimeric AapA1 forms size-specific pores, implying that the formed pores may have a specific pore size that allows passage of only molecules of a particular size or below.

We also found large fluctuations in the distance of K16/K23 residues from the center of the membrane (Figure 3D-F). To further understand the role of K16/K23 residues in AapA1-induced polar pore formation, we designed three constrained simulation systems and one mutation simulation system. Following constrained fixation of the K16/K23 residue side chains, the range of fluctuation of the AapA1 tilt angle in the transmembrane helical region became smaller, and the transport efficiency of water molecules through the transmembrane helical region was reduced by 16-fold (Figures 4A-J). No water molecules were translocated in CMD_hepta_K16/K23A simulations (Figure 4I-J). This finding suggests that the free-state movement of the K16/K23 residue side chains affects the backbone movement of AapA1, which is extremely important for the angle of insertion into the membrane and water permeability of the transmembrane barrel region; moreover, the movement of the K16/K23 residue side chains is critical for water molecule transfer (Figures 4A-J). In the calcein release assay, we also observed the impact of the K16/K23 mutants on water permeability (Figure 4K). However, we found that in the *in vitro* experiments, the K16/K23 mutations were insufficient to significantly reduce the water permeability function of the pores formed by AapA1, which differents from our simulation results to some extent (Figure 4). We propose two hypotheses: (1) The K16A/K23A double mutation in AapA1 protein may induce a shift in oligomerization states, favoring the formation of higher-order polymeric assembliesover the heptameric configuration; (2) Beyond the charged residues at positions 16 and 23, other conserved amino acids may critically modulate the intrinsic water permeability through steric gating or hydrogen-bonding network remodeling.

The AapA1 protein contains only positively charged and hydrophobic residues and no negatively charged residues. In general, the K11 residue always interacted electrostatically with the upper lipid membrane, and the K29 and R30 residues always bound to the lower lipid membrane (Figure 3A-C). However, the K16 and K23 residues within the transmembrane region did not interact with a particular layer of the lipid bilayer. We conducted WT-BEMetaD simulations to analyze the conformation states of the AapA1 monomeric protein in the membrane and found that the K16/K23 residue side chains can orient toward either the upper or lower layer of the membrane (Figure 5A). The three different conformation states were relatively stable, and the K16/K23 residue side chains were observed to oscillate between the upper and lower layers of the membrane (Figure 5B-C). The oscillation of the positively charged K16/K23 residue side chains could be an impetus for the accelerated transport of substances through the transmembrane pores.

Persister cells are a survival strategy of bacteria to resist killing by antibiotics by slowing down or halting the growth, reducing protein synthesis, and decreasing membrane potential ^60–63^. AapA1 expression in *H. pylori* leads to a rapid morphological transition from a helical bacterial cell to a coccobacillus form and then to a persister cell state. *H. pylori* in the dormant state shows some degree of clinical resistance to the combination of conventional therapy and vitamin D3 treatment ^9^. By conducting a vitamin D release assay, we found that multimeric AapA1-formed transmembrane pores allowed vitamin D3 permeation through the cell membrane and that this membrane permeability was controlled by amino acids at K16 and K23 (Figure 5D). The presence of AapA1-induced transmembrane pores may contribute to the exocytosis of vitamin D3, ergothioneine, etc., from the bacterial cell under unfavorable conditions such as antibiotic treatment and vitamin D3 co-treatment by altering membrane permeability, which subsequently transforms the growing bacterial cell into a dormant cell.

## Conclusions

The type I toxin protein AapA1-cell membrane system exhibits a high degree of specificity, which poses a challenge to study its sequence–structure–function relationship and mechanism of action through bioinformatics prediction tools and conventional experiments. Because MD simulations can capture subtle changes in the structure of chemical/biochemical components, we used MD simulations to probe the dynamic structure of AapA1-cell membrane systems and their mechanisms of action. AapA1 is involved in the formation of hydrophilic pores; this process is dependent on its selectivity for the cell membrane and the swing motion of the K16/K23 side chains. These findings suggest that the positively charged amino acid residues in AapA1 play a key role in facilitating the insertion of the polypeptide chain into the cell membrane and stabilizing the formation of hydrophilic channels.

## Acknowledgments

The study was funded by the National Natural Science Foundation of China (NSFC, grant numbers: 32171249 and 31670727), Natural Science Foundation of Shandong Province, China (ZR2024QC369) and the Research Foundation of Dezhou University (grant number: 2023xjrc208). We would like to thank TopEdit (www.topeditsci.com) for its linguistic assistance during the preparation of this manuscript.

## Supporting Information for

The lysine side chain’s swing motion in AapA1 toxin promotes water transport through transmembrane pores

**Fig. S1.**
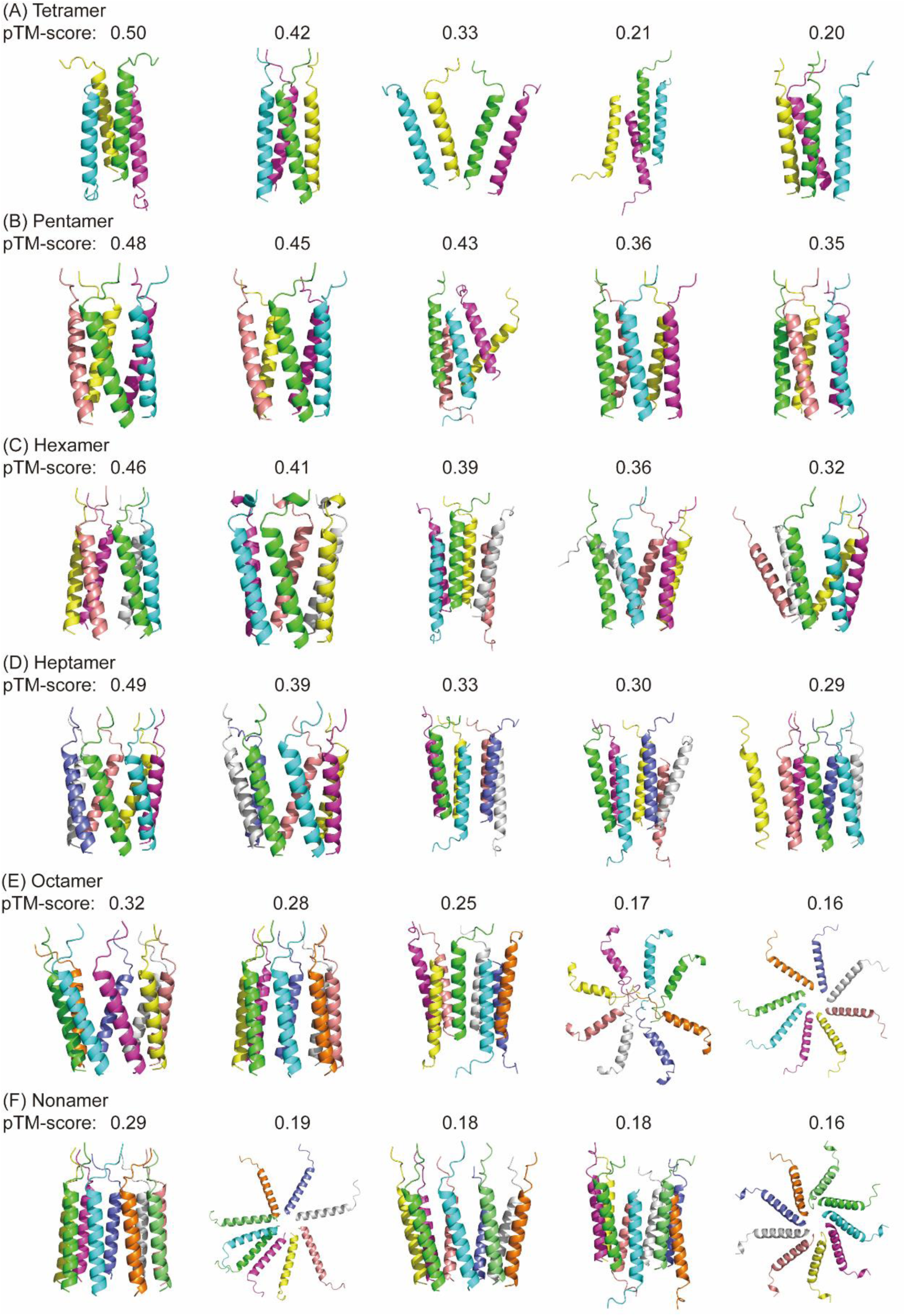
The top-5 predicted multimeric AapA1 toxin proteins structure and its corresponding pTM-score.

**Fig. S2.**
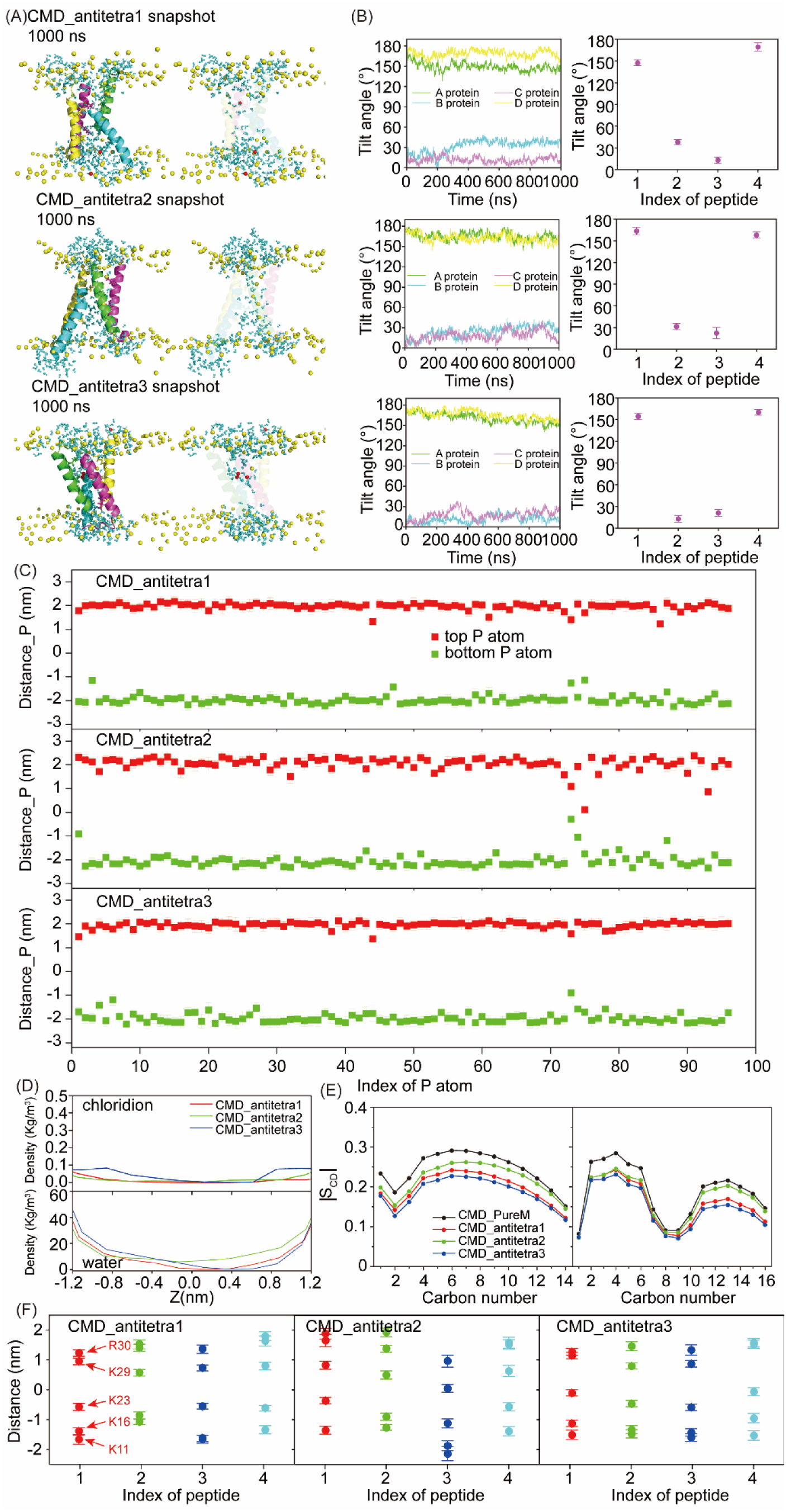
(A) The 1000th ns snapshots of the Antiparallel tetramer AapA1 and POPE/POPG (3:1) structure of CMD_antitetra simulations. In the snapshots, the following colors are used for the four AapA1 proteins: A protein, green; B protein, cyan; C protein, lightmagenta; D protein, yellow; the lipid phosphorus atoms are shown as yellow spheres, water are shown as cyan sticks, and chloridions are shown as red spheres. (B) The time series of fluctuation of tilt angles for each AapA1 protein (left) and the average tilt angles for each AapA1 protein during the last 200 ns (right). (C) The average distance of each phosphorus atom from the COM of the lipid bilayer during the last 200 ns of the trajectories in the CMD_antitetra simulations. “Distance_P” represents the average distance for phosphorus atoms. (D) Mass density of chloridions and water in the CMD_antitetra simulations. (E) Deuterium order parameters (|SCD|) of the saturated (left) and unsaturated (right) lipid acyl chains in the CMD_antitetra simulations. (F) The average distance of each COM of the Lys-11, Lys-16, Lys-23, Lys-29 and Arg-30 residue side chain from the COM of the lipid bilayer during the last 200 ns of the trajectories in the CMD_antitetra simulations.

**Fig. S3.**
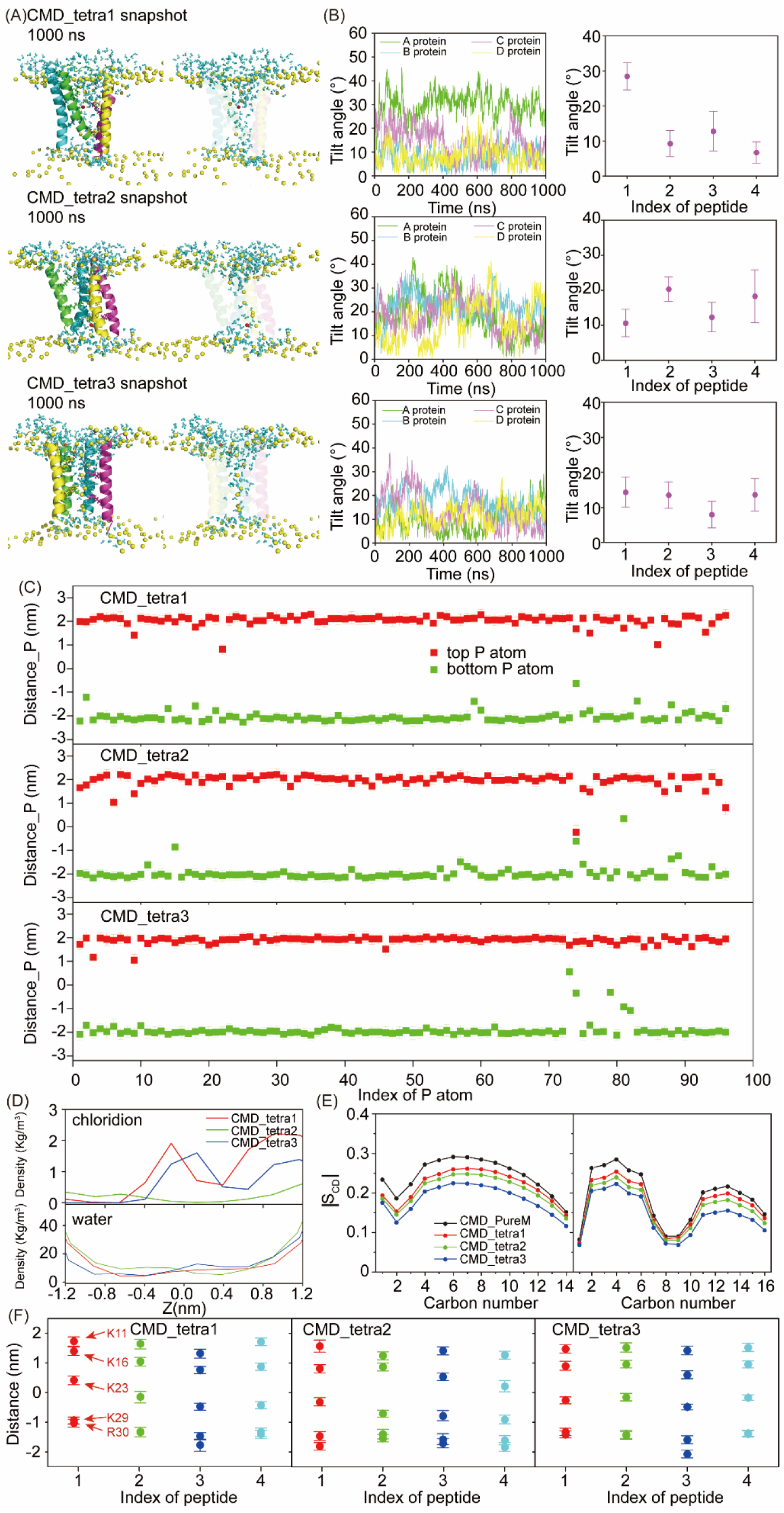
(A) The 1000th ns snapshots of the parallel tetramer AapA1 and POPE/POPG (3:1) structure of CMD_tetra simulations. In the snapshots, the following colors are used for the four AapA1 proteins: A protein, green; B protein, cyan; C protein, lightmagenta; D protein, yellow; the lipid phosphorus atoms are shown as yellow spheres, water are shown as cyan sticks, and chloridions are shown as red spheres. (B) The time series of fluctuation of tilt angles for each AapA1 protein (left) and the average tilt angles for each AapA1 protein during the last 200 ns (right). (C) The average distance of each phosphorus atom from the COM of the lipid bilayer during the last 200 ns of the trajectories in the CMD_tetra simulations. “Distance_P” represents the average distance for phosphorus atoms. (D) Mass density of chloridions and water in the CMD_tetra simulations. (E) Deuterium order parameters (|SCD|) of the saturated (left) and unsaturated (right) lipid acyl chains in the CMD_tetra simulations. (F) The average distance of each COM of the Lys-11, Lys-16, Lys-23, Lys-29 and Arg-30 residue side chain from the COM of the lipid bilayer during the last 200 ns of the trajectories in the CMD_tetra simulations.

**Fig. S4.**
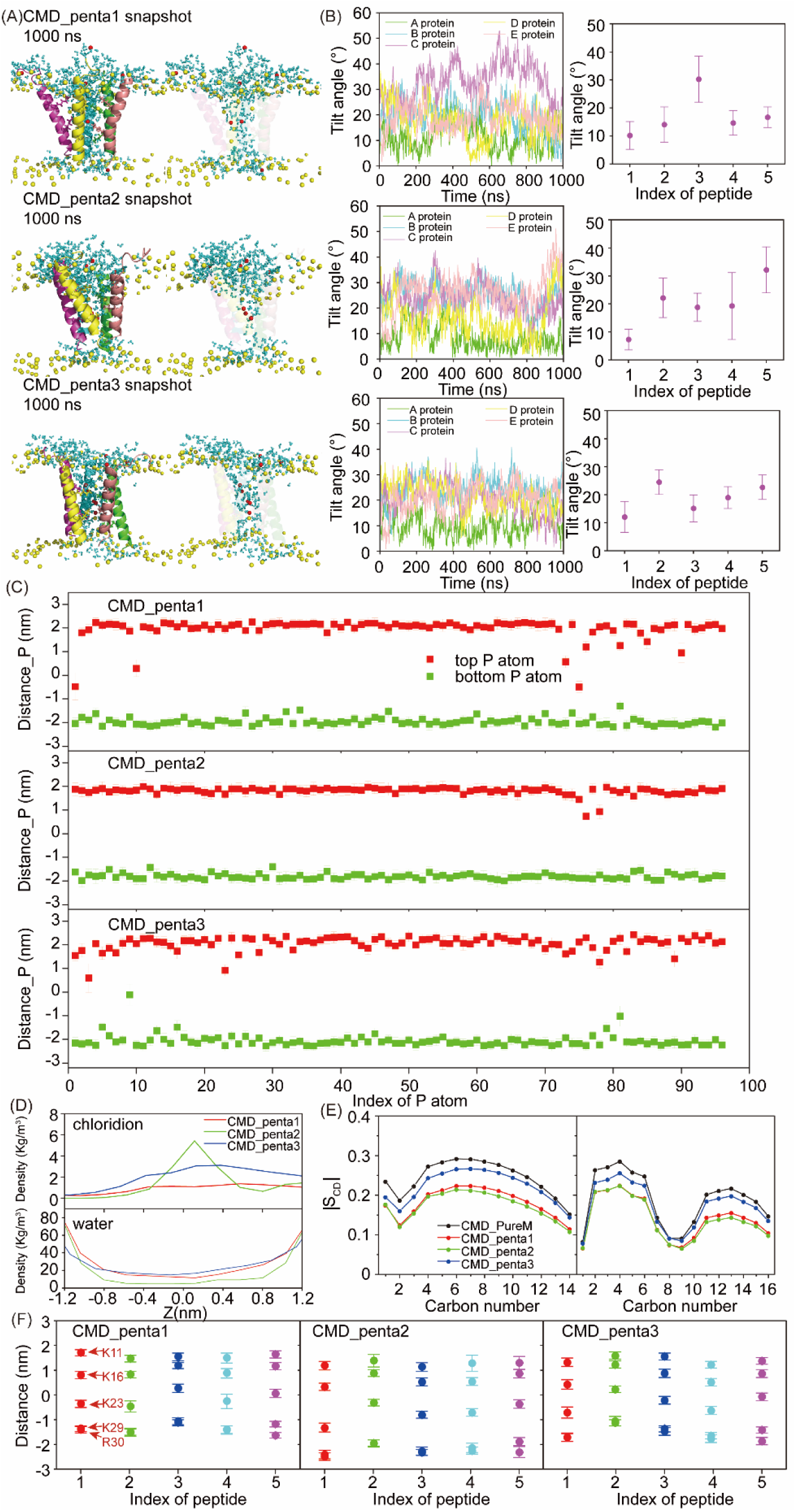
(A) The 1000th ns snapshots of the parallel pentamer AapA1 and POPE/POPG (3:1) structure of CMD_penta simulations. In the snapshots, the following colors are used for the five AapA1 proteins: A protein, green; B protein, cyan; C protein, lightmagenta; D protein, yellow; E protein, salmon; the lipid phosphorus atoms are shown as yellow spheres, water are shown as cyan sticks, and chloridions are shown as red spheres. (B) The time series of fluctuation of tilt angles for each AapA1 protein (left) and the average tilt angles for each AapA1 protein during the last 200 ns (right). (C) The average distance of each phosphorus atom from the COM of the lipid bilayer during the last 200 ns of the trajectories in the CMD_penta simulations. “Distance_P” represents the average distance for phosphorus atoms. (D) Mass density of chloridions and water in the CMD_penta simulations. (E) Deuterium order parameters (|SCD|) of the saturated (left) and unsaturated (right) lipid acyl chains in the CMD_penta simulations. (F) The average distance of each COM of the Lys-11, Lys-16, Lys-23, Lys-29 and Arg-30 residue side chain from the COM of the lipid bilayer during the last 200 ns of the trajectories in the CMD_penta simulations.

**Fig. S5.**
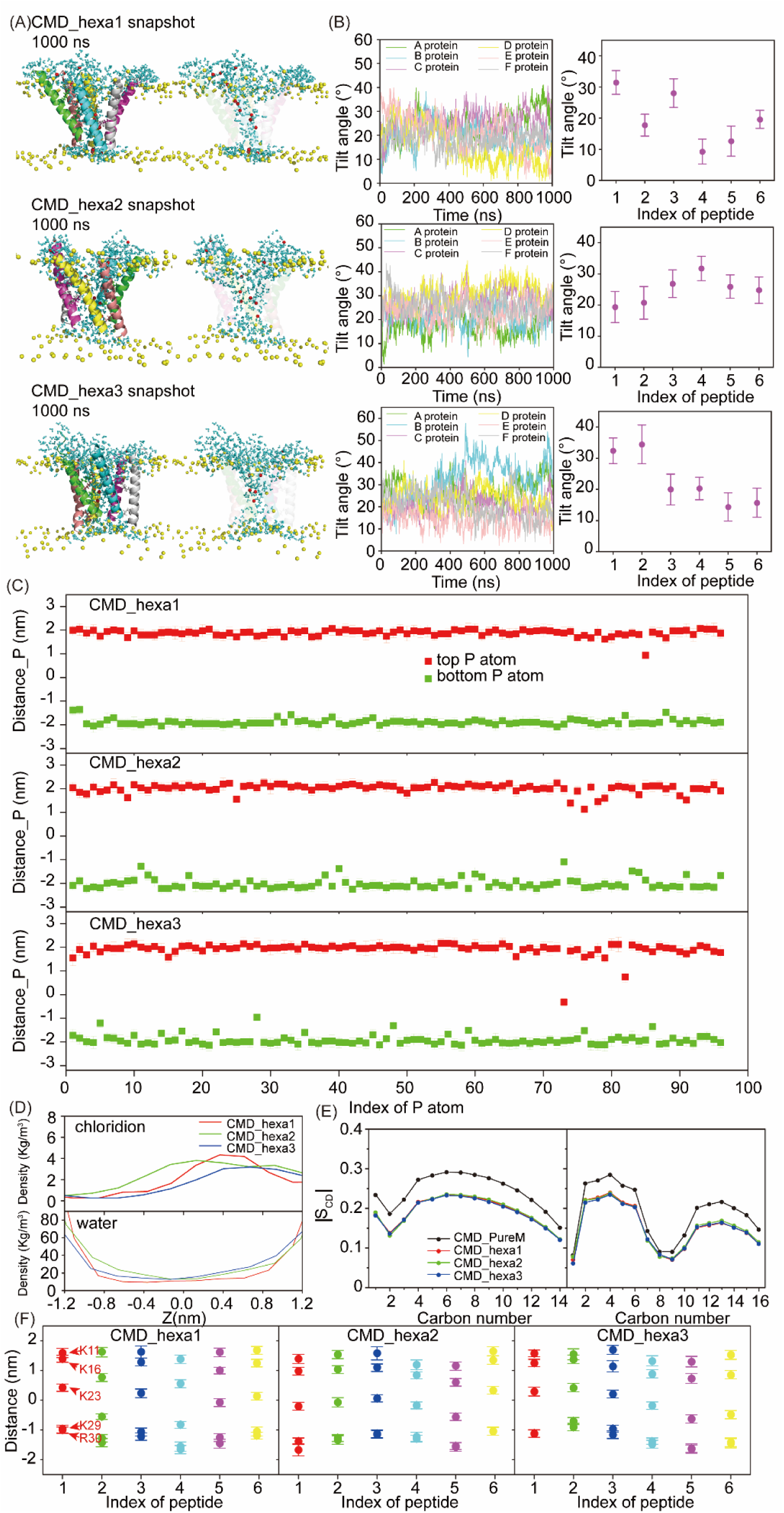
(A) The 1000th ns snapshots of the parallel hexamer AapA1 and POPE/POPG (3:1) structure of CMD_hexa simulations. In the snapshots, the following colors are used for the six AapA1 proteins: A protein, green; B protein, cyan; C protein, lightmagenta; D protein, yellow; E protein, salmon; F protein, gray; the lipid phosphorus atoms are shown as yellow spheres, water are shown as cyan sticks, and chloridions are shown as red spheres. (B) The time series of fluctuation of tilt angles for each AapA1 protein (left) and the average tilt angles for each AapA1 protein during the last 200 ns (right). (C) The average distance of each phosphorus atom from the COM of the lipid bilayer during the last 200 ns of the trajectories in the CMD_hexa simulations. “Distance_P” represents the average distance for phosphorus atoms. (D) Mass density of chloridions and water in the CMD_hexa simulations. (E) Deuterium order parameters (|SCD|) of the saturated (left) and unsaturated (right) lipid acyl chains in the CMD_hexa simulations.(F) The average distance of each COM of the Lys-11, Lys-16, Lys-23, Lys-29 and Arg-30 residue side chain from the COM of the lipid bilayer during the last 200 ns of the trajectories in the CMD_hexa simulations.

**Fig. S6.**
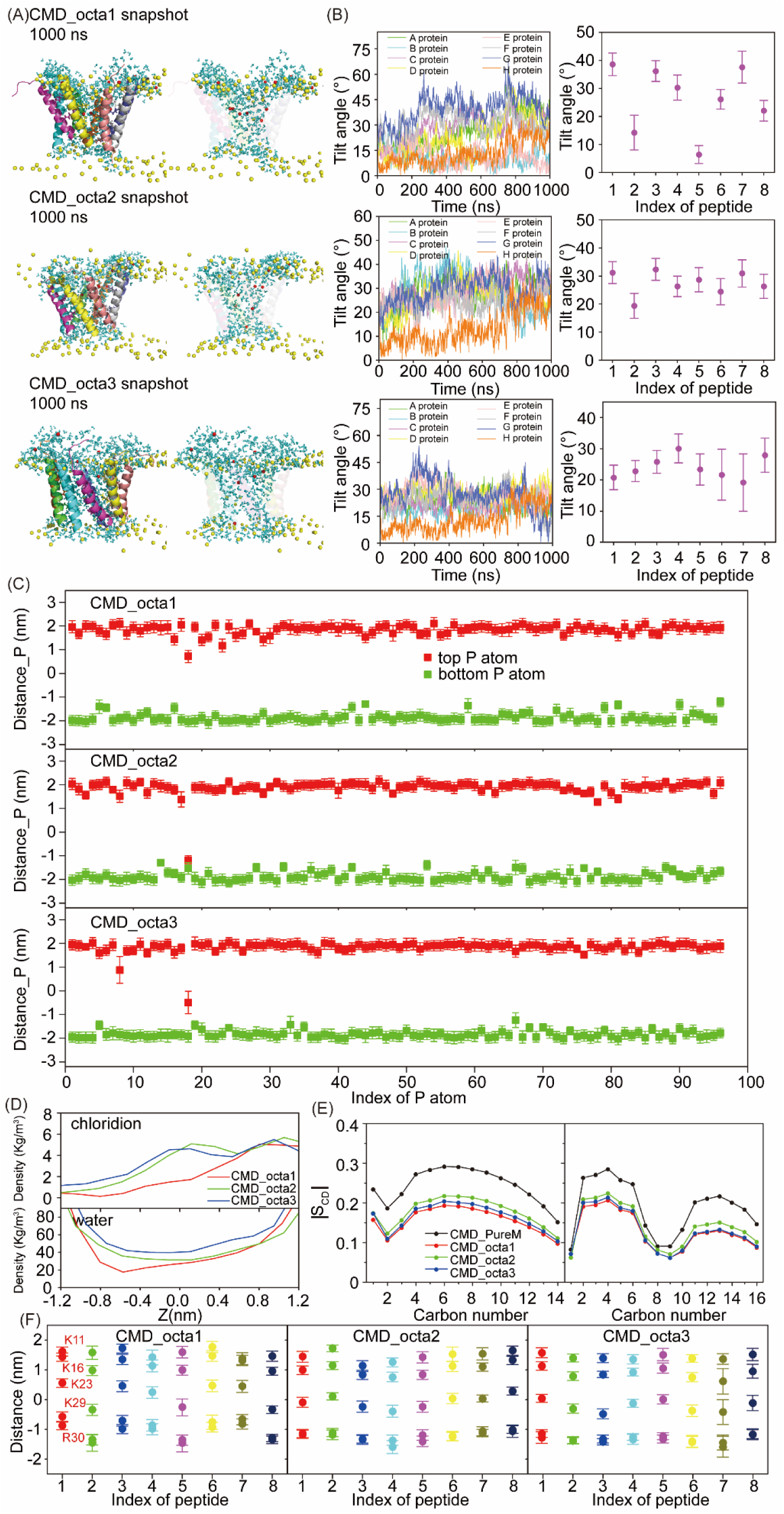
(A) The 1000th ns snapshots of the parallel octamer AapA1 and POPE/POPG (3:1) structure of CMD_octa simulations. In the snapshots, the following colors are used for the eight AapA1 proteins: A protein, green; B protein, cyan; C protein, lightmagenta; D protein, yellow; E protein, salmon; F protein, gray90; G protein, slate; H protein, orange; the lipid phosphorus atoms are shown as yellow spheres, water are shown as cyan sticks, and chloridions are shown as red spheres. (B) The time series of fluctuation of tilt angles for each AapA1 protein (left) and the average tilt angles for each AapA1 protein during the last 200 ns (right). (C) The average distance of each phosphorus atom from the COM of the lipid bilayer during the last 200 ns of the trajectories in the CMD_octa simulations. “Distance_P” represents the average distance for phosphorus atoms. (D) Mass density of chloridions and water in the CMD_octa simulations. (E) Deuterium order parameters (|SCD|) of the saturated (left) and unsaturated (right) lipid acyl chains in the CMD_octa simulations. (F) The average distance of each COM of the Lys-11, Lys-16, Lys-23, Lys-29 and Arg-30 residue side chain from the COM of the lipid bilayer during the last 200 ns of the trajectories in the CMD_octa simulations.

**Fig. S7.**
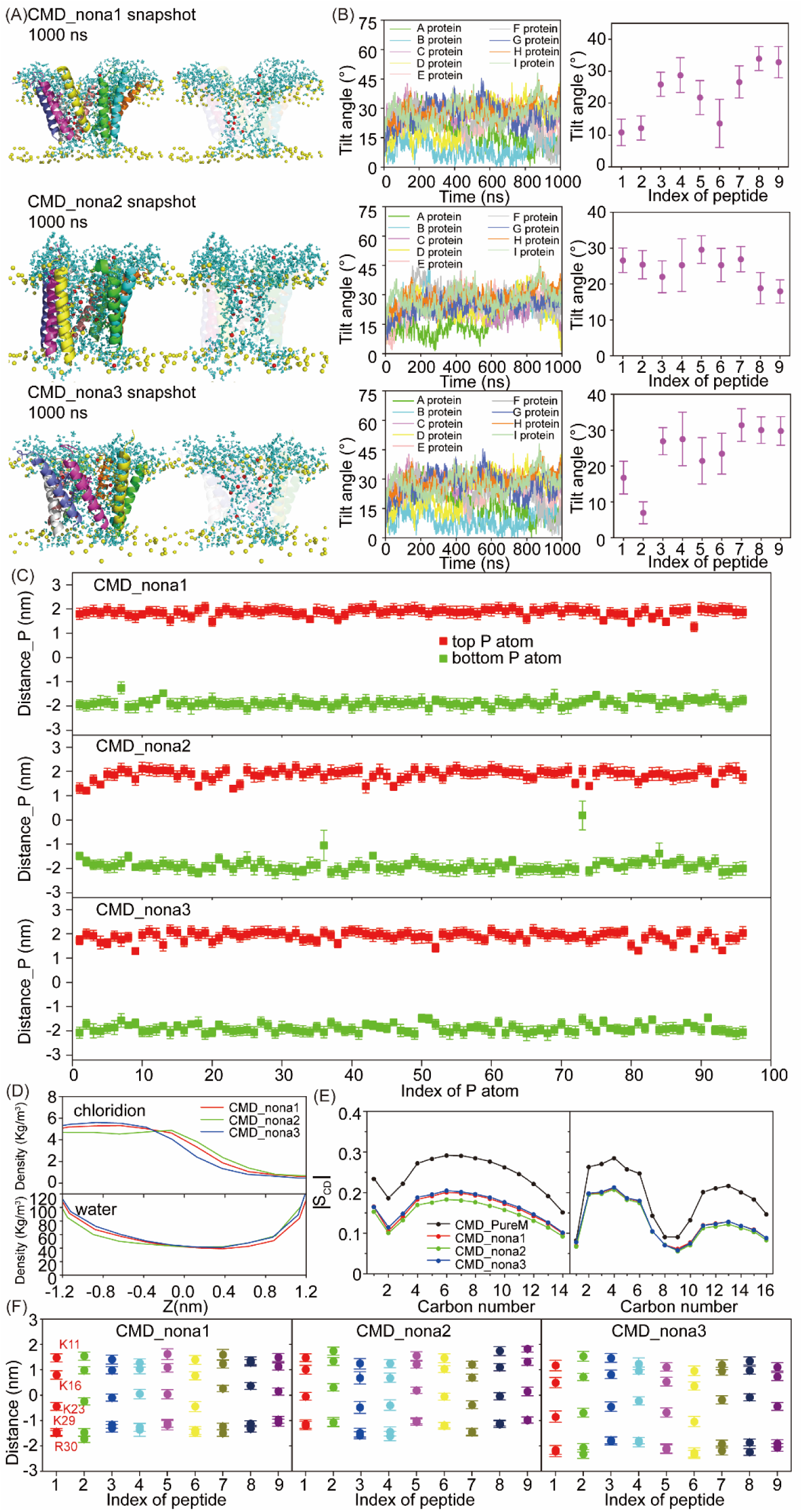
(A) The 1000th ns snapshots of the parallel nonamer AapA1 and POPE/POPG (3:1) structure of CMD_ nona simulations. In the snapshots, the following colors are used for the eight AapA1 proteins: A protein, green; B protein, cyan; C protein, lightmagenta; D protein, yellow; E protein, salmon; F protein, gray90; G protein, slate; H protein, orange; I protein, lime; the lipid phosphorus atoms are shown as yellow spheres, water are shown as cyan sticks, and chloridions are shown as red spheres. (B) The time series of fluctuation of tilt angles for each AapA1 protein (left) and the average tilt angles for each AapA1 protein during the last 200 ns (right). (C) The average distance of each phosphorus atom from the COM of the lipid bilayer during the last 200 ns of the trajectories in the CMD_ nona simulations. “Distance_P” represents the average distance for phosphorus atoms. (D) Mass density of chloridions and water in the CMD_ nona simulations. (E) Deuterium order parameters (|SCD|) of the saturated (left) and unsaturated (right) lipid acyl chains in the CMD_ nona simulations. (F) The average distance of each COM of the K11, K16, K23, K29 and R30 residue side chain from the COM of the lipid bilayer during the last 200 ns of the trajectories in the CMD_nona simulations.

**Fig. S8.**
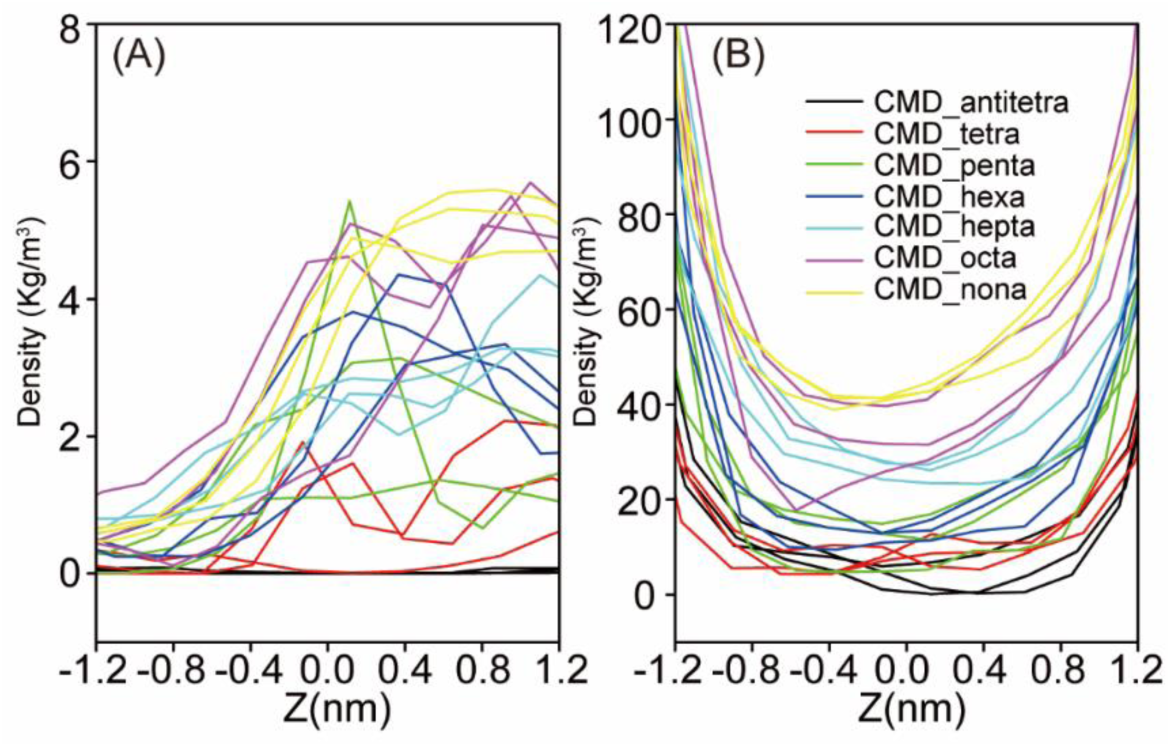
Mass density of chloridions (A) and water (B) in seven different multimeric AapA1 proteins simulations.

**Fig. S9.**
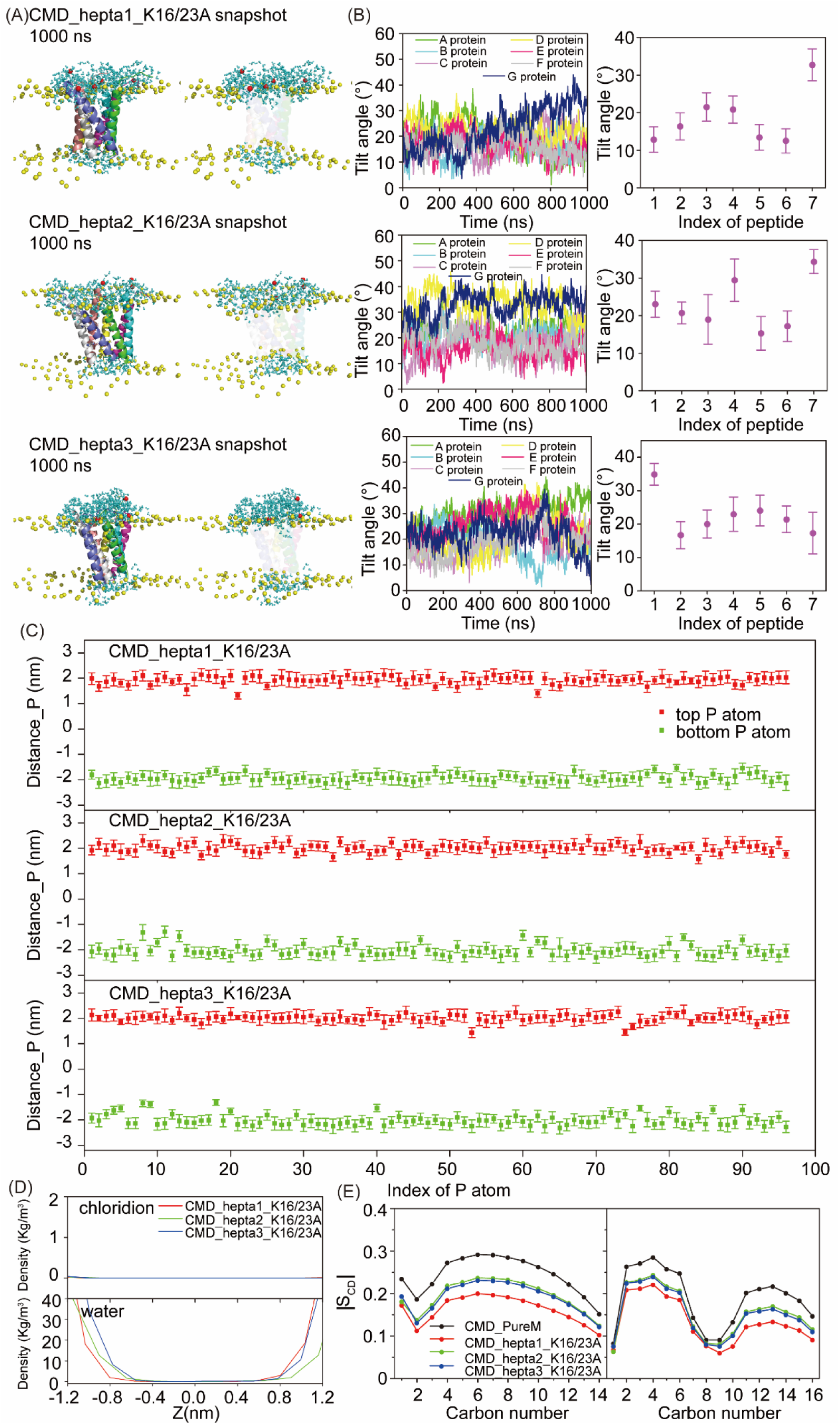
(A) The 1000th ns snapshots of the Heptamer AapA1 with K16/23A mutation and POPE/POPG (3:1) structure of CMD_hepta_K16/23A simulations. In the snapshots, the following colors are used for the seven AapA1 proteins: A protein, green; B protein, cyan; C protein, lightmagenta; D protein, yellow; E protein, salmon; F protein, gray90; G protein, slate; the lipid phosphorus atoms are shown as yellow spheres, water are shown as cyan sticks, and chloridions are shown as red spheres. (B) The time series of fluctuation of tilt angles for each AapA1 K16/23A protein (left) and the average tilt angles for each AapA1 K16/23A protein during the last 200 ns (right). (C) The average distance of each phosphorus atom from the COM of the lipid bilayer during the last 200 ns of the trajectories in the CMD_hepta_K16/23A simulations. “Distance_P” represents the average distance for phosphorus atoms. (D) Mass density of chloridions and water in the CMD_hepta_K16/23A simulations. (E) Deuterium order parameters (|SCD|) of the saturated (left) and unsaturated (right) lipid acyl chains in the CMD_hepta_K16/23A simulations.

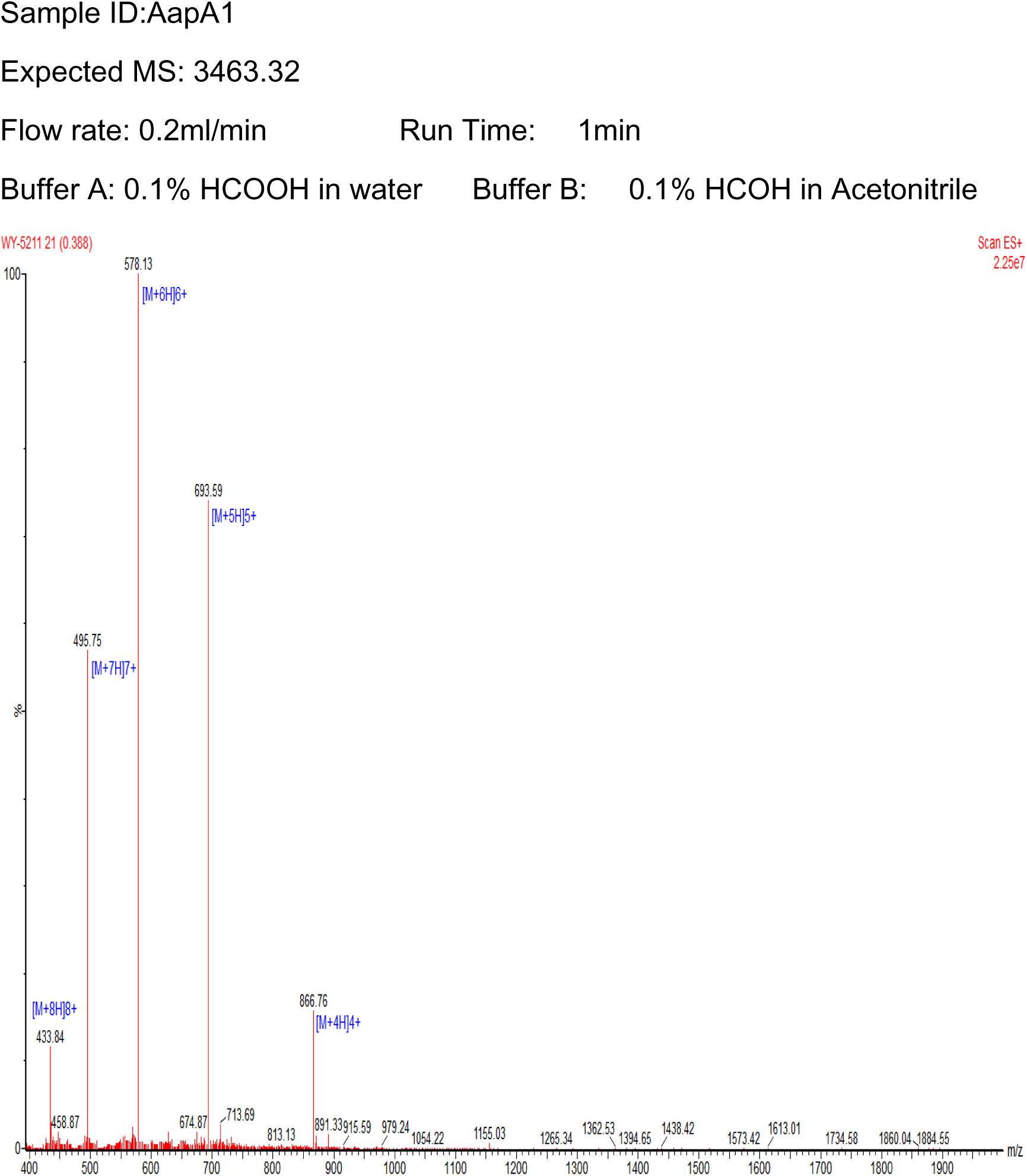

